# Molecular phenotyping of colorectal neoplasia shows dynamic and adaptive cancer stem cell population admixture

**DOI:** 10.1101/2022.06.11.495729

**Authors:** Ester Gil Vazquez, Nadia Nasreddin, Gabriel N Valbuena, Eoghan J Mulholland, Hayley L Belnoue-Davis, Holly Eggington, Ryan O Schenck, Valérie M Wouters, Pratyaksha Wirapati, Kathryn Gilroy, Tamsin R M Lannagan, Dustin J Flanagan, Arafath K Najumudeen, Sulochana Omwenga, Amy M B McCorry, Alistair Easton, Viktor H Koelzer, James E East, Dion Morton, Livio Trusolino, Timothy Maughan, Andrew D Campbell, Maurice B Loughrey, Philip D Dunne, Petros Tsantoulis, David J Huels, Sabine Tejpar, Owen Sansom, Simon J Leedham

**Author notes:** Correspondence should be addressed to S.J.L. Joint first authors (Ester Gil Vazquez, Nadia Nasreddin).

## Abstract

Intestinal homeostasis is underpinned by *LGR5+ve* crypt-base columnar stem cells (CBCs), but following injury, dedifferentiation results in the emergence of *LGR5-ve* regenerative stem cell populations (RSCs), characterised by fetal transcriptional profiles. Neoplasia hijacks regenerative signalling, so we assessed the distribution of CBCs and RSCs in mouse and human intestinal tumors. Using combined molecular-morphological analysis we demonstrate variable expression of stem cell markers across a range of lesions. The degree of CBC-RSC admixture was associated with both epithelial mutation and microenvironmental signalling disruption, and could be mapped across disease molecular subtypes. The CBC-RSC equilibrium was adaptive, with a dynamic response to acute selective pressure, and adaptability was associated with chemoresistance. We propose a fitness landscape model where individual tumors have equilibrated stem cell population distributions along a CBC-RSC phenotypic axis. Cellular plasticity is represented by position shift along this axis, and is influenced by cell-intrinsic, extrinsic and therapeutic selective pressures.

## Introduction

The intestine is an exemplar of an adult stem cell-supported tissue system. The identification of selective expression of *Leucine-rich repeat-containing G-protein coupled receptor 5 (Lgr5)* in the crypt-base columnar cells (CBC) of both human and murine crypts enabled lineage tracing and the demonstration of the first *bona fide* intestinal stem cell marker (Barker *et al*., 2007). Although *Lgr5+ve* CBCs demonstrably underpin steady state intestinal homeostasis, their immediate contribution in supporting epithelial regeneration is less clear. *Lgr5* expression declines to undetectable following colitis induction in murine models (Davidson *et al*., 2012) and recovers at day 5 post injury, suggesting that alternative stem cell population(s) support the early response to epithelial damage. Recent work has shown dedifferentiation and adaptive reprogramming of multiple cell types within residual epithelium (de Sousa and de Sauvage, 2019), with reversion to a primitive molecular phenotype and induction of fetal intestinal gene expression through activation of epithelial Yap signalling (Nusse *et al*., 2018; Yui *et al*., 2018). Individual genes within this fetal transcriptional programme, such as *Ly6a (Sca1), Anxa1* (Nusse *et al*., 2018; Yui *et al*., 2018) and *Clu* (Ayyaz *et al*., 2019) have been used to identify regenerative cells with stem cell properties, (collectively termed regenerative stem cells, RSCs, from here on).

Cancer is often described as the ‘wound that never heals’ through co-option and corruption of physiological cell signalling. Neoplasia is characterised by the activity of stem cells, from the impact of initiating (epi)mutation through to the emergence of therapy resistant clones and metastatic seeding (Fumagalli *et al*., 2020). In mouse models, genetic inactivation of the key CRC driver gene, *Adenomatous Polyposis Coli* (*Apc*) in *Lgr5*+ve cells precipitated rapid tumour induction, confirming CBCs as a cell-of-origin in intestinal tumorigenesis (Barker *et al*., 2009). However, murine studies subsequently showed that induction of inflammation and disruption of homeostatic morphogen gradients could result in neoplasia originating from *Lgr5*-ve cells, outside of the crypt base (Schwitalla *et al*., 2013; Davis *et al*., 2015). Furthermore, selective and effective killing of *Lgr5* cells had no impact on primary tumour growth (de Sousa e Melo *et al*., 2017) and the migratory cells that seed and colonise distant organs were frequently *Lgr5*-ve at dissemination (Fumagalli *et al*., 2020). In humans, recent integrated analysis of single cell data demonstrated that serrated polyps arise from differentiated cells through a gastric metaplastic process (Chen *et al*., 2021), and a proportion of established colorectal tumours have minimal expression of *LGR5* (Merlos-Suarez *et al*., 2011; Morral *et al*., 2020; Shimokawa *et al*., 2017). Elegant recent work has shown that both subpopulations of *Lgr5*+ve and -ve tumor cells have elevated rDNA transcription and protein synthesis characteristic of functional stem cell activity (Morral *et al*., 2020), and that lineage conversion between cell types can be driven by the combination of key CRC driver genes and microenvironmental extracellular signalling (Han *et al*., 2020). Together these data indicate (1) the presence of alternative/additional (*Lgr5*-ve) stem cell populations in neoplastic lesions, and (2) that induced cell plasticity allows primary tumors to adapt to the loss of individual cancer stem cell populations.

Natural selection acts upon phenotype, so the capacity to easily measure a definable and pathologically-relevant cancer molecular phenotype that can temporally track cancer cell fate is central to the concept of assessing tumor evolutionary trajectory. Stem cell plasticity underpins intestinal regeneration, and it is evident that *Lgr5* cannot be used as a sole marker for putative cancer stem cell populations in established lesions. Here we have undertaken molecular and morphological analysis to assess the stem cell molecular phenotype across a range of mouse neoplasia models, derived organoids and human lesions, and examine the factors that influence phenotypic plasticity. We propose a conceptual phenotypic fitness landscape model to contextualise the relationship between neoplastic stem cell activity (fitness) and cellular phenotype. *Lgr5*+ve crypt-base columnar and *Lgr5*-ve regenerative stem cells represent distinct but interlinked fitness peaks along a stem cell phenotypic axis. In individual untreated tumours, there is a distribution of stem cell phenotypes along this axis that reaches an equilibrium point, determined by combination of selected epithelial mutation and microenvironmental signalling. Phenotypic plasticity can be represented by a shifting in the phenotype distribution of the stem cell population, is regulated by emergent adaptive signalling pathways, and is required for tumor adaptation to therapeutic selective pressures.

## Results

### Application of molecular signatures and morphological markers to identify intestinal stem cell populations

First, we selected an established crypt-base columnar cell signature (Munoz *et al*., 2012) and defined an RSC signature by aggregating published regenerative stem cell signatures (Mustata *et al*., 2013; Yui *et al*., 2018) and refining these based on information on cell type-associated expression from single cell RNAseq data (Methods). Next, we used fluorescent *in situ* hybridisation, multiplex immunohistochemistry and human single cell RNA expression data to assess normal cell compartment expression of CBC and RSC genes (Figure 1A-C). In steady state, *LGR5* expression (CBC cell marker) was seen in discrete cell populations and was confined to epithelial cells at the base of the crypts. In contrast, we saw no homeostatic epithelial expression of *ANXA1* (a widely used regenerative stem cell marker) (Nusse *et al*., 2018; Yui *et al*., 2018) in normal human colon or murine small intestine, although low level expression was seen in the distal colon of mice. However, *ANXA1* expression was detected in stromal, myeloid and T-cells both in human scRNA datasets and on-slide in mouse and human tissue. (Figure 1A,B). Additional regenerative morphological stem cell markers, *Ly6a* (mouse) and *PLAUR* (human as there is no human ortholog of *Ly6a*), were also assessed across all lesions (Figure S1).

**Figure 1.**
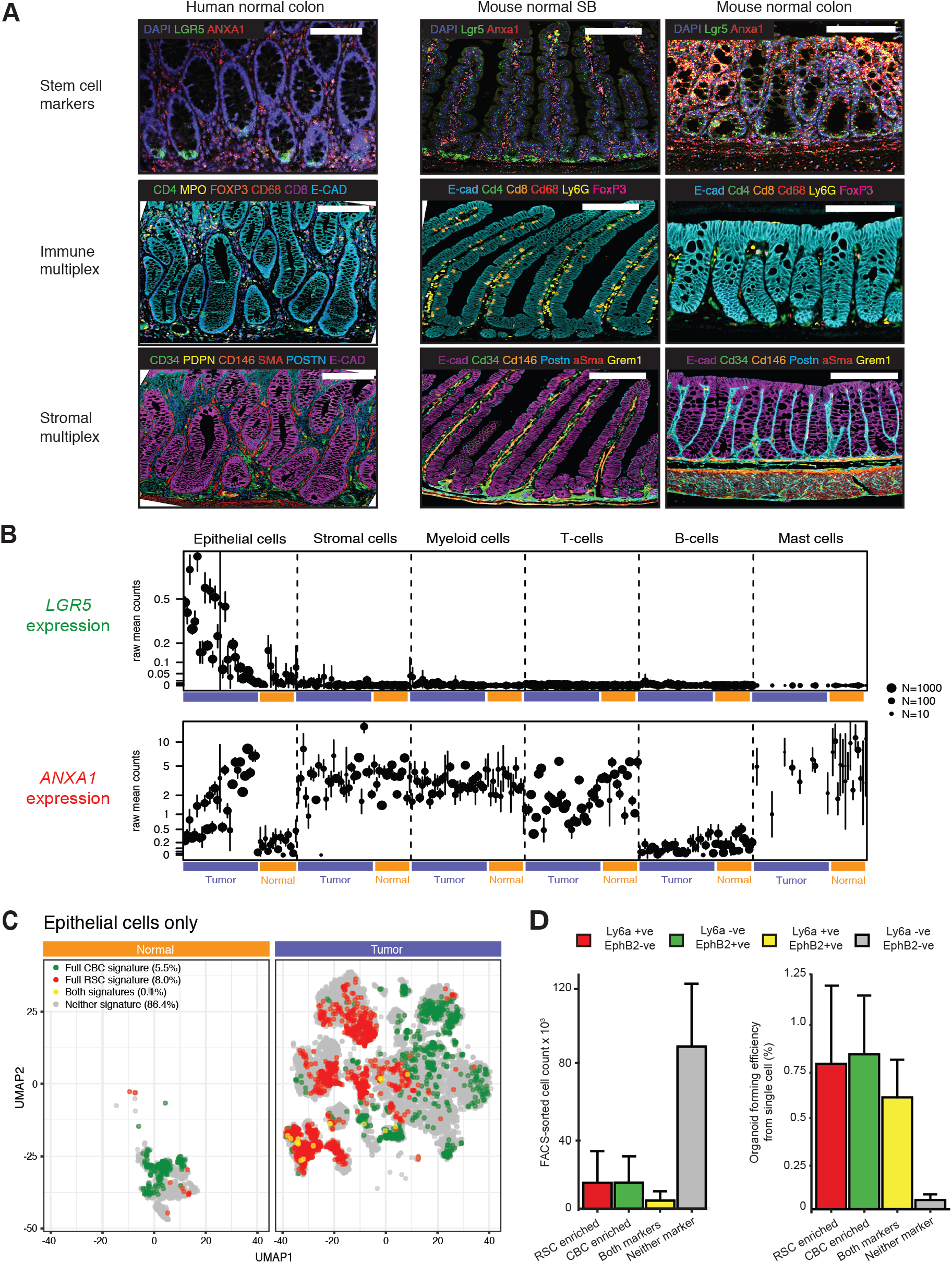
Molecular and morphological assessment of the crypt-base columnar and regenerative stem cell marker expression spectrum. **A**. Dual color ISH and multiplex IHC to show expression pattern of representative CBC and RSC cell markers alongside homeostatic immune and stromal cell distribution in normal mouse and human intestine. Scale bar 100μm **B**. Distribution of human multicompartmental scRNA expression of *LGR5* (CBC marker) and *ANXA1* (RSC marker) in normal (orange bars) and cancer cell compartments (purple bars). **C**. UMAP plot of single epithelial cells from human normal and colorectal cancer samples showing cell populations enriched for CBC (green cells), RSC (red cells) and mixed CBC and RSC gene expression (yellow cells). Cells with no enriched stem cell signature expression are grey. **D**. Stem cell marker expressing cell count and organoid forming efficiency from plated single cells following FACS segregation of KPN mouse primary tumours, measured at day 7 post seeding.

Assessment of cell-specific expression of *LGR5* and *ANXA1* from scRNA of human cancers revealed an inversely proportional enrichment of these individual cell markers in the cancer epithelial cell compartment, indicating that individual tumors may be variably skewed towards CBC- or RSC-predominant stem cell phenotypes (Figure 1B). Discrete stem cell populations could also be distinguished by application of full CBC and RSC gene profiles in human normal tissue and cancer scRNA datasets, with a small number of cells expressing both stem cell signatures (Figure 1C). Together these data show that despite the cell compartment promiscuity of some regenerative stem cell markers, application of CBC and RSC expression signatures and morphological markers can be used to identify different epithelial stem cell populations in mouse and human intestinal tissue.

### Stem cell potential of mouse tumour CBC and RSC enriched cell populations

To demonstrate that cells expressing CBC and RSC cell markers retain stem cell potential in tumors and to test whether CBC and RSC occupy comparable fitness peaks, we turned to mouse models of intestinal carcinogenesis to undertake *ex vivo* single cell clonogenicity experiments. New mouse models cumulatively combine multiple alleles to replicate key CRC epithelial driver gene mutations, and generate a range of disease states that phenocopy human polyposis syndromes and the CRC consensus molecular subtypes (Jackstadt *et al*., 2019). Intestinal cancers from the autochthonous *Vil1-CreER*^*T2*^*;Kras*^*G12D*^*;p53*^*fl/fl*^*;Rosa26*^*N1ICD/+*^ (KPN) line were digested and cell suspensions subjected to established protocols for cell segregation and organoid generation, previously used to enrich for CBCs (Merlos-Suarez *et al*., 2011) and RSCs (Yui *et al*., 2018) from mouse models. Controlled flow gating strategies (Figure S2) across individual tumors obtained variable numbers of CBC and RSC enriched cell populations for each biological repeat, but consistent with human scRNA data (Figure 1C), there were numerically fewer intermediate cells expressing both sets of stem cell surface markers (Figure 1D). Single cell organoid generation experiments demonstrated that cell populations enriched for CBC, RSC or both cell markers were capable of single cell organoid generation, indicating retained stem cell potential, whereas cells without expression of any stem cell marker had very little clonogenic capacity *ex vivo* (Figure 1D).

### Stem cell marker expression varies across a range of human tumor molecular subtypes and mouse intestinal neoplasia models

Next, we assessed the phenotypic landscape in bulk transcriptome samples from CRC, to see if we could infer neoplastic stem cell population admixture using these translationally relevant sample sets. We calculated single sample enrichment scores for our molecular CBC and RSC molecular signatures using gene set variation analysis (GSVA, Hänzelmann *et al*., 2013) and derived an intestinal stem cell index measurement by subtracting the CBC score from the RSC score. This allowed assessment of the relative abundance of different stem cell signatures across a wide range of human and mouse tumors.

First, we applied the stem cell index and morphological markers to human polyp and CRC bulk transcriptome data from the S:CORT dataset. In both polyps and tumors, we observed variation in expression of stem cell molecular signatures and morphological markers along a phenotypic axis, indicating variable enrichment/admixture of CBC and RSC stem cell phenotypes. Individual lesions clustered by histological or molecular subtype across this axis, with CBC predominance in conventional pathway lesions such as tubulovillous adenomas, CMS2 and CRIS-C,D,E cancer subtypes. Conversely, relative enrichment for RSC was seen in serrated precursor lesions and tumor molecular subtypes CMS4 and CRIS-B (Figure 2A-C). *In situ* hybridisation confirmed differential epithelial marker *LGR5* and *ANXA1/PLAUR* expression across representative lesions (Figure 2D,E, and S1).

**Figure 2.**
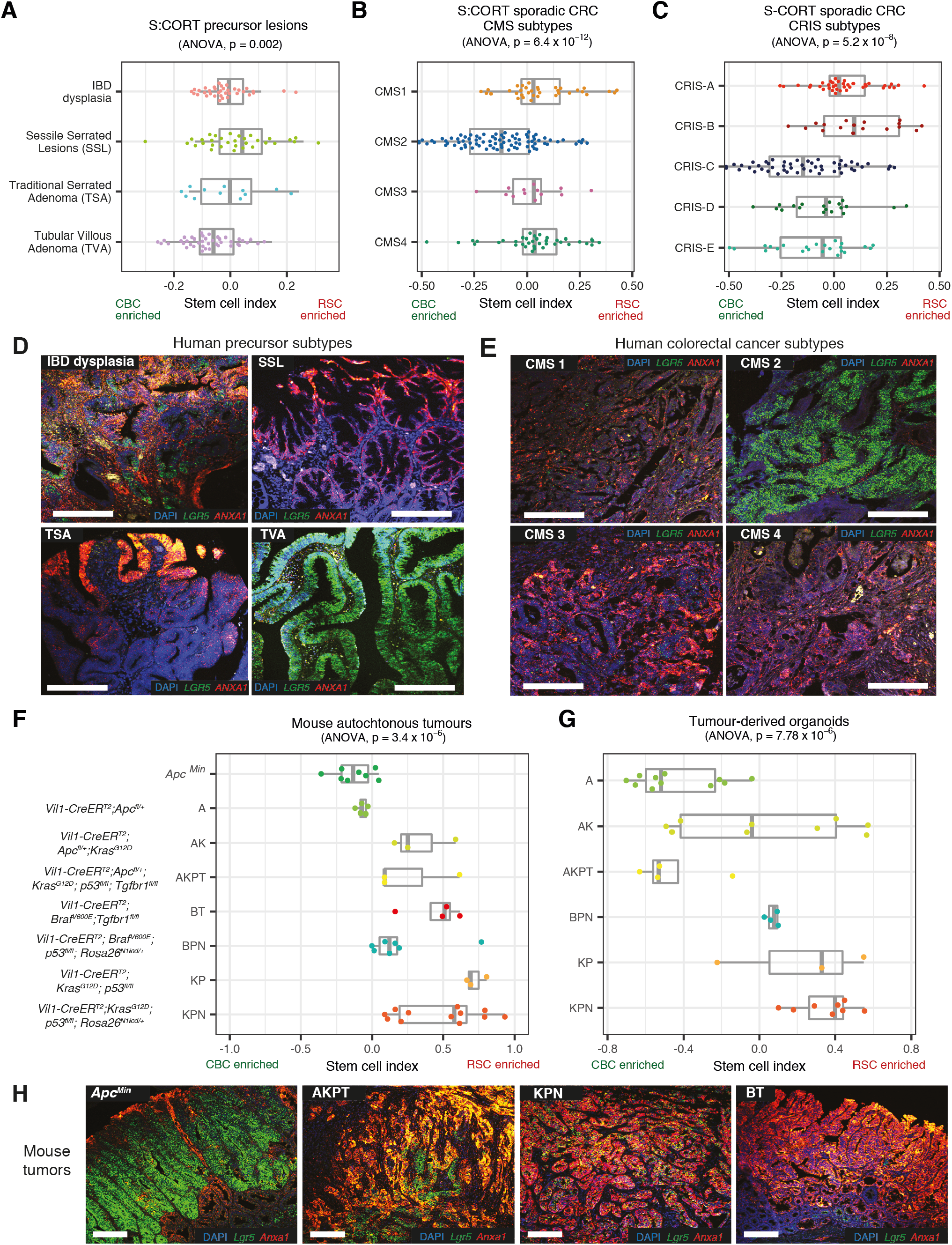
Application of stem cell index to mouse and human neoplasia. Using stem cell index to map: **A**. human colorectal precursor lesions **B**. colorectal cancer consensus molecular subtypes (CMS) and **C**. colorectal cancer intrinsic subtypes (CRIS) across a molecularly defined CBC to RSC expression spectrum (using S-CORT datasets). **D-E**. Dual color ISH for *LGR5* (CBC marker - green) and *ANXA1* (RSC marker - red) expression in **D**. representative human precursor lesions and **E**. representative human colorectal cancers segregated by consensus molecular subtype. **F**. Using stem cell index to map mouse autochthonous tumors, and, **G**. matched derived organoids, across a molecularly defined CBC to RSC expression spectrum. **H**. Dual color ISH for *Lgr5* (CBC marker - green) and *Anxa1* (RSC marker - red) expression in representative genotype tumors across the CBC to RSC spectrum. Statistical analysis, ANOVA, p values as stated. All animals crossed with *Vil-CreER*^*T2*^. Scale bars 100μm. Driver alleles initialisation - A is *Apc*^*fl/+*^, *Apc*^*Min*^ is *Apc*^*Min*^, B is *Braf*^*V600E*^, K is *Kras*^*G12D*^, P is *p53*^*fl/fl*^, T is *Tgfβr1*^*fl/fl*^, N is *Rosa26*^*N1icd/+*^

Next, we applied the mouse version of the stem cell index to murine bulk transcriptome data. Mouse models permit control over the epithelial genotype, which allowed assessment of the cumulative impact of individual driver genes on molecular phenotype in both autochthonous tumors and derived organoids. Similar to human lesions, we observed variation in the stem cell index across a CBC to RSC phenotypic axis, with individual lesions clustering by mouse epithelial genotype (Figure 2F). Transgenic manipulation of some key drivers produced a notable skew in cell admixture. Thus, epithelial *Apc* mutation enriched for CBC cell marker expression, whereas Mapk activation (through *Kras* or *Braf* mutation) or Tgfβ disruption (through *Tgfβr1* knockout) skewed the stem cell index towards RSC markers. In multi-allele models, the cumulative accumulation of driver genes correlated with variable degrees of admixture of stem cell marker expression. In mouse cancer organoids, the baseline stem cell index was distinct for each genotype and the pattern of stem cell index distribution across different genotypes was conserved between the *ex-vivo* bulk tumour transcriptome and the derived *in vitro* organoids, cultured in the absence of other niche cellular constituents (Figure 2G). This indicates the predominant contribution of changes in epithelial gene expression to the variance in the stem cell index. Multicolour *in situ* hybridisation was used to confirm variable epithelial expression of CBC and RSC stem cell markers in four key disease positioned animal models: *Apc*^*Min*^, AKPT, KPN, and BT (Figure 2H). These models were selected to represent conventional and serrated molecular carcinogenesis pathways and span the stem cell phenotypic axis (Jackstadt *et al*., 2019; Lannagan *et al*., 2021; Leach *et al*., 2021). Together these data demonstrate that the stem cell index can be used to assess variable admixture of stem cell phenotypes across a range of mouse and human tumors.

### Driver genes and pathways associated with variable stem cell molecular phenotype

To assess human tumour genotype-stem cell phenotype correlations, we looked for associations of key driver gene mutations with stem cell phenotype in published human single cell datasets (Lee *et al*., 2020). We found significant correlation between epithelial *APC* and *BRAF* mutations with single cell transcriptome-derived CBC and RSC phenotypes respectively. However, there was no association with other key drivers such as *p53* and *KRAS* which were seen in cells across the phenotypic spectrum (Figure S3A). Next, we mapped the distribution of the stem cell index in TCGA lesions stratified by key CRC driver mutations to see if the stem cell index variation mirrored that seen with the clearly defined mouse genotypes (Figure 3A). We then segregated TCGA tumours into polarised deciles for stem cell marker expression (Figure 3C) and undertook pairwise comparison of single nucleotide variation (Figure S3B) and copy number variation (Figure S3C) between CBC and RSC predominant tumours. In the tumors most enriched for CBC, ligand-independent wnt mutations (*APC* and *CTNNB1*) were found in 84% of tumors but only 35% of RSC predominant lesions, where ligand-dependent wnt alterations such *RNF43, ZNRF3*, and *RSPO2/3* fusions were also seen (Figure 3B). There was also variation in the mutation prevalence and type impacting the MAPK/PI3K pathways and the TGFβ superfamily (Figure 3B). Together, these data indicate an association between optimally selected driver gene mutations in key signalling pathways and the predominant stem cell phenotype in human lesions.

**Figure 3.**
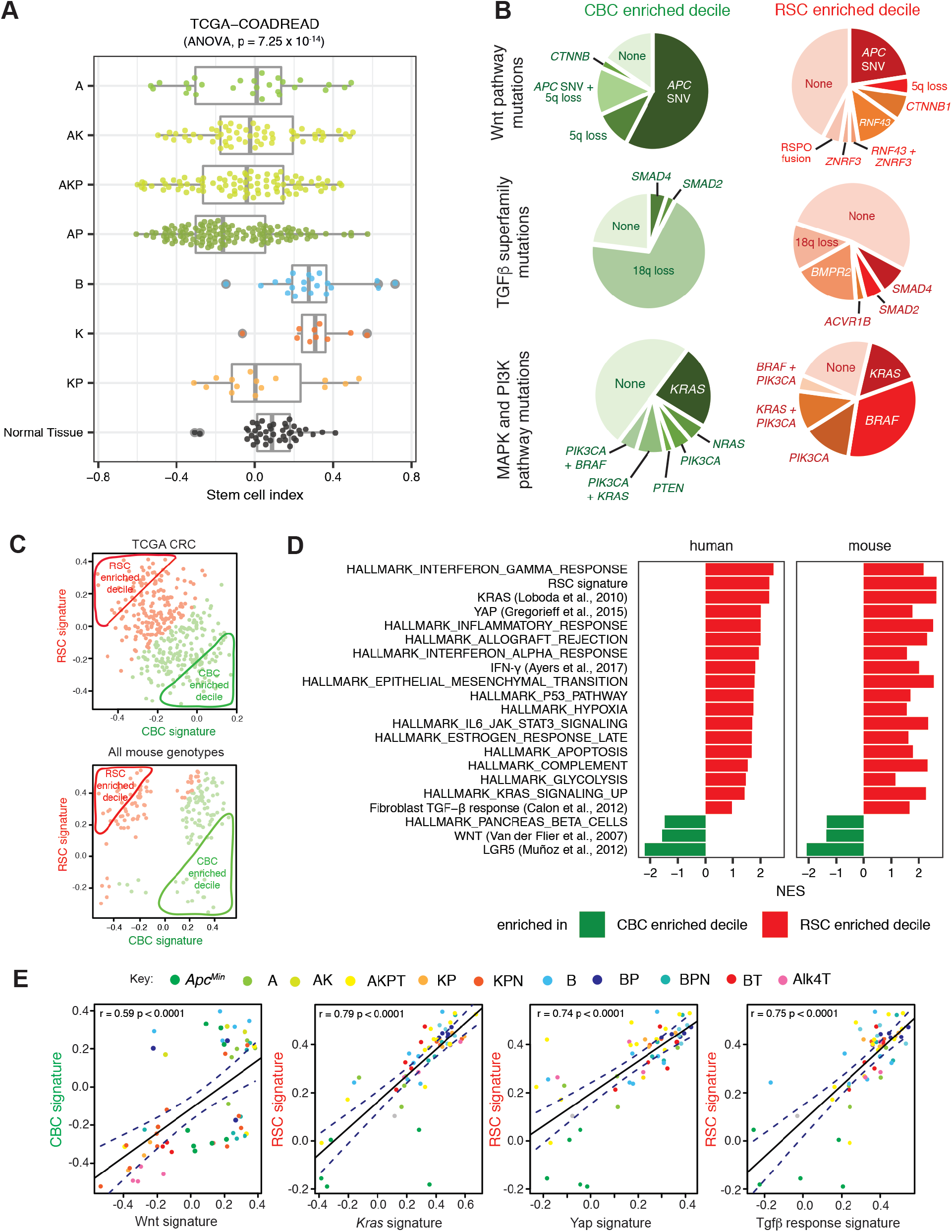
Driver genes and pathways associated with variable stem cell molecular phenotype. **A**. Human genotype-stem cell phenotype correlation based on stem cell index distribution in TCGA tumors with different putative driver gene SNV mutation genotypes, contrasted to normal tissue from same dataset (driver gene initials: A is APC, K is KRAS, P is p53, B is BRAF). **B**. Comparison of mutation type and prevalence disrupting the Wnt pathway, MAPK and PIK3CA pathways and the TGFβ superfamily in TCGA tumors subdivided into CBC and RSC predominant deciles. **C**. Segregation of mouse and human lesions by CBC (x-axis) and RSC (y-axis) signature expression. Predominant (above median) expression signature in each tumor defined by color (CBC in green, and RSC in red), and the 10% most polarised CBC or RSC expressing tumors were segregated into CBC and RSC enriched deciles for comparison **D**. Gene set enrichment analysis of hallmark and select pathways in bulk transcriptome from human tumours (TCGA) and murine lesions (Glasgow dataset) segregated into CBC and RSC predominant deciles. Pathways shown have *P*_FDR_ ≤0.25 apart from YAP in the mouse lesions and Fibroblast TGF-β response in the human tumors. **E**. Correlation of key pathway expression signatures with CBC or RSC gene expression across a range of mouse models. Different genotypes identified by different colors as determined by the key. Driver alleles initialisation - A is *Apc*^*fl/+*^, *Apc*^*Min*^ is *Apc*^*Min*^, B is *Braf*^*V600E*^, K is *Kras*^*G12D*^, P is *p53*^*fl/fl*^, T is *Tgfβr1*^*fl/fl*^, N is *Rosa26*^*N1icd/+*^, Alk4 is *Alk4* ^*fl/fl*^.

Next, we used transcriptome data from the TCGA and our suite of animal models to compare cross-species signalling pathway disruption. Mouse and human tumours were segregated by stem cell index, and gene set enrichment analysis used to contrast hallmark pathway disruption in the most CBC- and RSC-predominant deciles. Strikingly, we saw significant cross-species correlation between stem cell phenotype and signalling pathway disruption, with activation of wnt signalling in CBC-predominant lesions, and enrichment of KRAS, YAP, TGFβ, and inflammatory pathways (such as IFN-γ) in RSC-enriched mouse and human tumors (Figure 3C,D). In mouse models, we mapped expression of these key pathways with CBC or RSC gene signatures across a large range of defined genotypes (Figure 3E). Although we identify some key cross-species candidate signalling hubs, we also show that there are a number of genotype-specific pathways associated with a predominant CBC or RSC phenotype across three of our key mouse genotypes (*Apc*^*Min*^, KPN and AKPT) (Figure S3D). Thus, it seems likely that individual tumours utilise both shared, and genotype/tumor-specific cell-intrinsic and extrinsic pathways to establish phenotypically convergent stem cell populations.

Together these data imply that epithelial ligand-independent Wnt signalling mutation (such as APC) enhances fitness of the crypt base columnar stem cell phenotype, whereas regenerative stem cell fitness may be influenced by signalling disruption from both epithelial cell-intrinsic (e.g KRAS, BRAF) and tumor microenvironmental sources, with some key pro-regenerative stem cell pathways mapping predominantly to immune (IFN-γ), stromal (TGFβ) and matrix (YAP) cell compartments. In light of this observation, we used multiplex staining to assess the cell compartment landscape of representative mouse tumors from our four key genotypes selected to span across the CBC to RSC stem cell phenotypic axis. We found quantifiable differences in the immune, stromal and matrix landscapes in tumors from different models (Figure 4A,B), with the matrix compartment, in particular, showing both interesting intra-tumour topographic heterogeneity (Figure S4A) and inter-tumour diversity between lesions from different genotypes (assessed using the Shannon index, Figure S4B). This was consistent with differential landscaping of the tumour context in the lesions from each of the different models, generating variable microenvironmental niches. We hypothesised that niche crosstalk back to the epithelium (through secreted signalling or mechano-transduction) could influence epithelial stem cell phenotype. To test the effect of secreted signalling directly, we turned to mouse and human tumour organoids and used cytokine/morphogen supplementation of the media to model the influence of immune and stromal cell signalling respectively. We saw a significant shift in the stem cell index towards an RSC enriched phenotype in normal organoids following media supplementation of both IFN-γ and TGFβ. In mouse cancer organoids, the genotype-specific stem cell phenotype was not fixed: we saw similar directional shifts in the stem cell index in the response of KPN mouse cancer organoids to both IFN-γ and TGFβ. However, AKPT organoids only responded to IFN-γ, consistent with the knockout of the Tgfβr1 receptor in this genotype. This shows that manipulation of immune- and stromally-derived signalling pathways can directly modulate epithelial phenotype. Furthermore, although the baseline stem cell phenotypic state is defined by organoid genotype (Figure 2G), it is not completely fixed by cancer cell driver mutation alone, and remains at least partly responsive to microenvironmental signalling.

**Figure 4.**
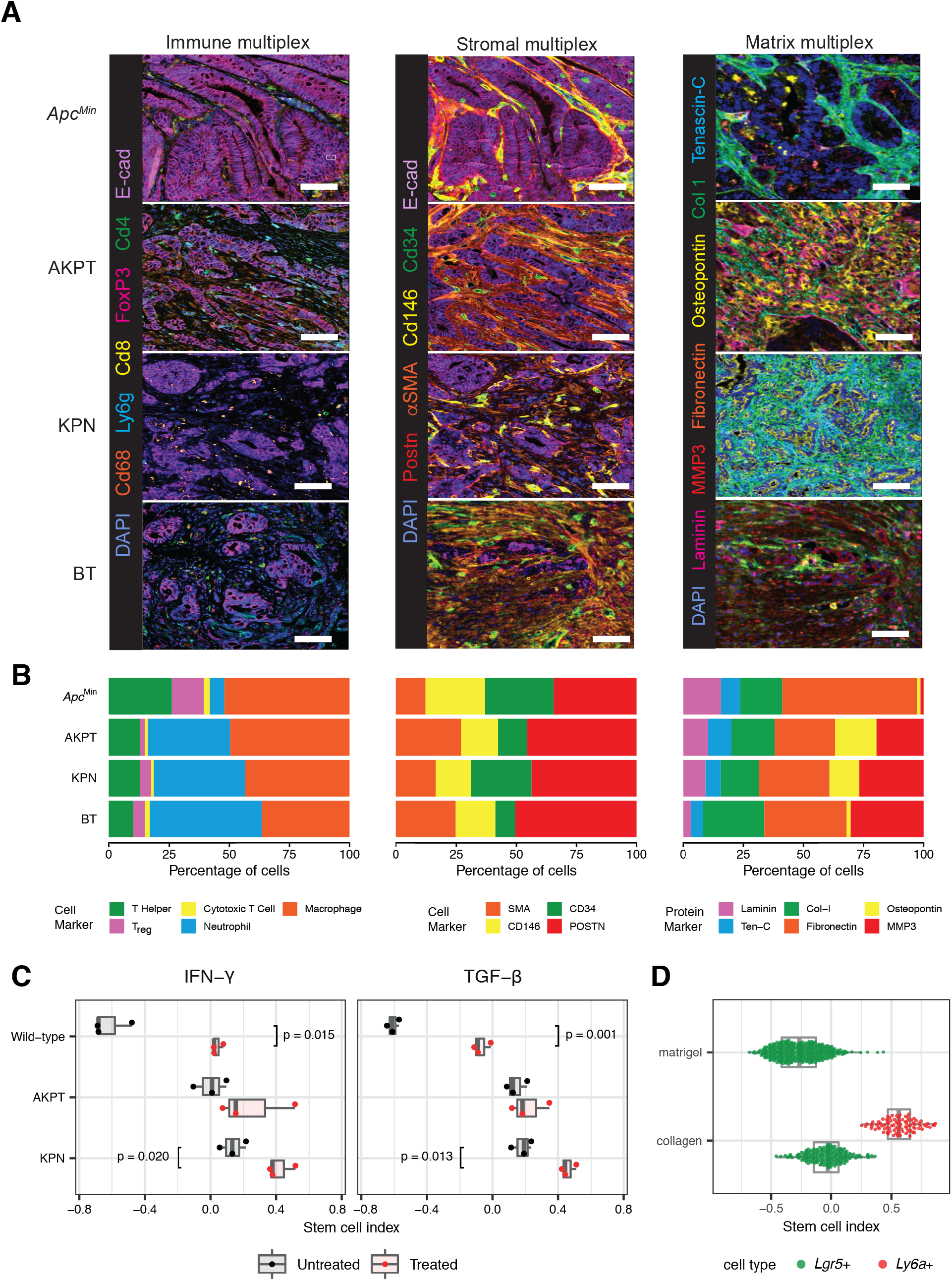
Microenvironmental landscaping and crosstalk influences epithelial stem cell phenotype. **A**. Representative multiplex IHC images of immune, stromal and matrix landscapes in mouse tumors selected from across the stem cell phenotypic axis. Scale bars 100μm. **B**. Variable proportion of different cell/matrix components in tumors from each genotype quantified from multiplex IHC images (n=3 mice per genotype). **C**. Impact of media supplementation of IFNγ (1μl/ml) and TGFβ1 (0.5μl/ml) on stem cell phenotype of wildtype and AKPT, KPN mouse tumour organoids. Ttest, p values as stated. **D**. Stem cell index applied to single cell transcriptome data derived from organoids grown in matrigel or collagen matrix (from Ramadan *et al*., 2021).

Constitutive YAP activation, through Lats1/2 kinase knockout induces fetal signature gene expression in murine organoids (Cheung *et al*., 2020). We (Koppens *et al*., 2021; Ramadan *et al*., 2021) and others (Yui *et al*., 2018) have shown that organoid growth in collagen can induce YAP and model the impact of matrix mechano-transduction (Figure S4C-E). To assess the effect of YAP activation on stem cell phenotype, we analysed single cell data from organoids grown in collagen and matrigel from Ramadan *et al*. (2021). We see the emergence of a Ly6a+ cell population only in organoids maintained in collagen, driving a significant shift in the stem cell index towards the regenerative phenotype (Figure 4D). This shift was confirmed in two independent bulk organoid transcriptome datasets, E-MTAB-5247 (Yui *et al*., 2018, Figure S4D) and E-MTAB-10082 (Ramadan *et al*., 2021, Figure S4E).

Together these data demonstrate that variance in the stem cell molecular phenotype is associated with differences in epithelial cell-intrinsic driver gene mutations alongside conserved cross-species disruption of key microenvironmental signalling pathways. Intercompartmental crosstalk from variably landscaped immune, stromal and matrix compartments across different tumours can directly influence epithelial stem cell phenotype through secreted signalling or mechano-transduction pathways.

### A dynamic equilibrium exists between neoplastic stem cell marker expressing populations

From our molecular analysis of large tumour sets, it was evident that expression of stem cell markers ranged across a CBC to RSC phenotypic axis, with single cell assessment, FACS segregation and morphological assessment of different lesions all demonstrating the co-existence of both discrete CBC, RSC and shared marker-expressing cell populations in mouse and human tumors. Following injury, regenerative stem cells are capable of reconstituting lost CBC cell populations (reviewed in de Sousa and de Sauvage, 2019), so we reasoned that the dysregulated signalling of the tumor milieu permitted a dynamic stem cell equilibrium with adaptive interconversion between cells situated on CBC and RSC stem phenotype peaks. To test this *in vitro*, we identified a dose of media IFN-γ that induced the most significant shift in stem cell phenotype in organoids grown in matrigel (Figure 5A). We then cultured Lgr5-GFP mouse organoids to permit direct flow cytometric detection of an *Lgr5*+ve stem cell population, and used high-dose media IFN-γ to apply a selective pressure to GFP-labelled cells. Flow cytometry analysis showed that IFN-γ media supplementation rapidly skewed cell phenotype, with all organoid cells including both GFP-high expressing CBC cells and GFP-low expressing progenitor cells, profoundly upregulating *Ly6a* expression consistent with a plastic adaptive shift along the CBC to RSC stem cell phenotypic axis (Figure 5B).

**Figure 5.**
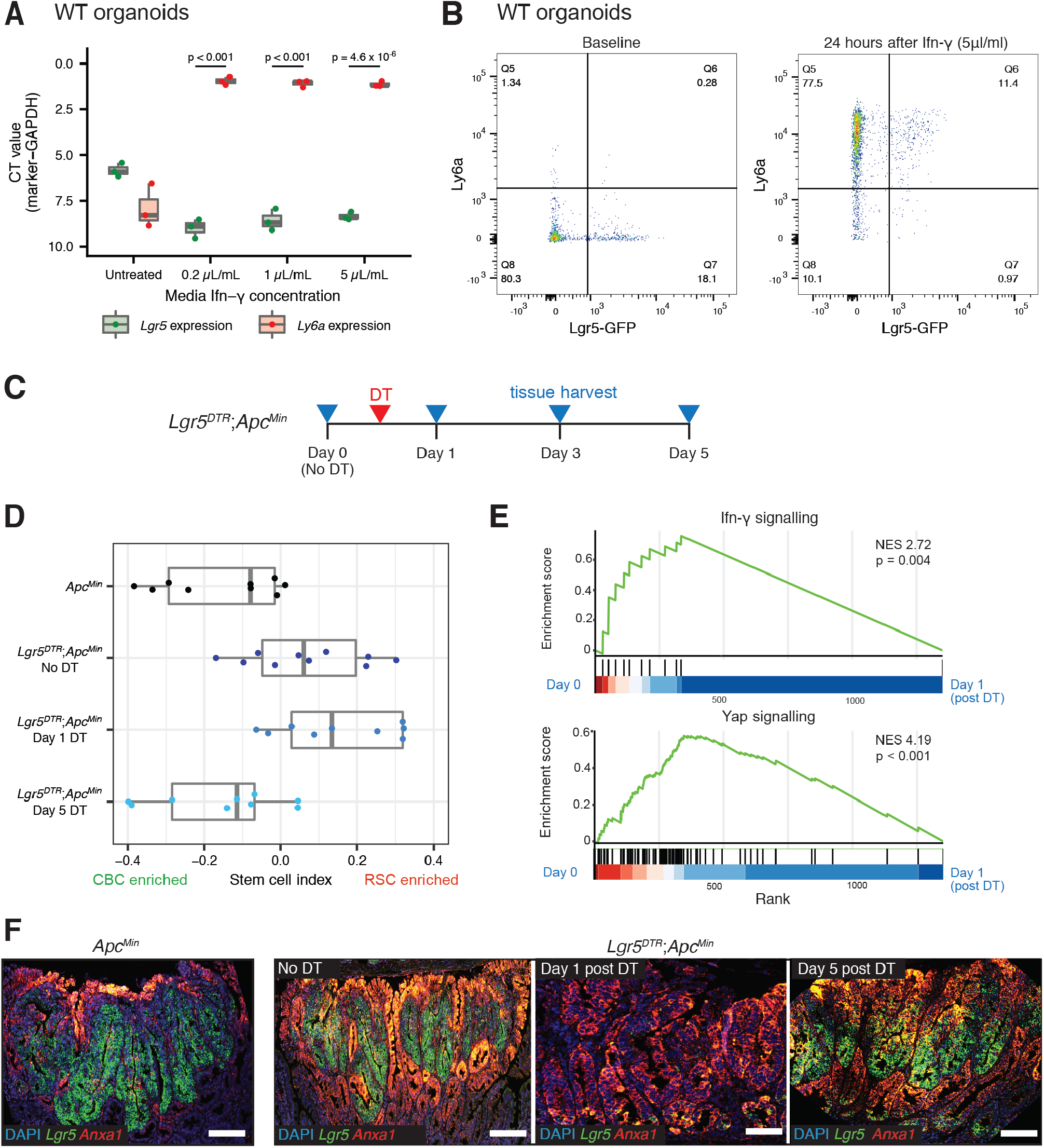
Adaptive shift of stem cell phenotype under selective pressure. **A**. Shift in stem cell marker expression detected by qRT-PCR following exposure of wild-type organoids to increasing concentrations of media IFN-γ. Statistical analysis, t-test, p values as stated. **B**. Skewed expression of stem cell markers, detected by FACS for Ly6a and GFP, following exposure of Lgr5-GFP labelled murine organoids to 5μl/ml of media IFN-γ. **C**. Schematic showing timing of recombination, diphtheria toxin activation and tissue harvesting of *Lgr5*^*DTR*^*;Apc*^*Min*^ mice. **D**. Using stem cell index to map polyp tissue from *Apc*^*Min*^ and *Lgr5*^*DTR*^*;Apc*^*Min*^ to show dynamic change in stem cell molecular phenotype measured by stem cell index, before and after DTR activation and CBC cell ablation **E**. Gene set enrichment analysis showing enrichment of Ifn-γ signalling and Yap signalling between Day 0 (unrecombined) and Day 1 (after DTR stem cell ablation). **F**. Dual color ISH for *Lgr5* (CBC marker green) and *Anxa1* (RSC marker red) to show marker expression change in *Apc*^*Min*^ and *Lgr5*^*DTR*^*;Apc*^*Min*^ polyps before and after CBC cell ablation.

To test whether this adaptive response to selective pressure could occur *in vivo*, we crossed *Apc*^*Min*^ animals with the *Lgr5*^*DTR*^ allele established by Tian *et al*. (2011) to generate *Lgr5*^*DTR*^*;Apc*^*Min*^ animals. As previously noted (Tian *et al*., 2021), targeted introduction of the DTR cassette and *Lgr5* hemizygosity impacted *Lgr5*+ve CBC number in *Lgr5*^*DTR*^*;Apc*^*Min*^ mouse polyps at steady state (Figure S5A) and in polyps (Figure S5B) and resulted in a detectable shift in the stem cell index in comparison with *Apc*^*Min*^ lesions alone (Figure 5D). Following diptheria toxin injection, almost all *Lgr5*+ve cells in established *Lgr5*^*DTR*^*;Apc*^*Min*^ mouse polyps were selectively ablated and we tracked the molecular phenotypic response to acute loss of the predominant stem cell population (Figure 5D-F). After 24 hours, ablation of *Lgr5*+ve cells provoked a dramatic upregulation of RSC in *Lgr5*^*DTR*^*;Apc*^*Min*^ polyps, followed by partial reconstitution and recovery of *Lgr5* expressing CBC by day 5 (Figure 5D,F). These shifts in the stem cell molecular phenotype were associated with acute upregulation of Ifn-γ, Yap (Figure 5E) and Kras signalling pathways (Figure S5C) but notably, there was no significant impact on polyp size, cell proliferation or apoptotic cell death following recombination (Figure S5E,F). This indicates that following selective ablation, rapid adaptive shifts in tumor signalling act to rapidly restore the stem cell equilibrium and that these profound shifts in stem cell phenotype can occur without detectable change in conventional tumor clinical response measurements.

### Temporally spaced assessment of stem cell index can be used to assess adaptive response to selective pressure

As part of a fitness landscape model, we propose that interlinked CBC and RSC stem population peaks co-exist in CRC, and that in untreated tumors, different combinations of epithelial mutation and microenvironmental signalling shift the stem cell population distribution to an equilibrium set point that varies between individual lesions. Given that there is some clustering of individual lesion set points within the established human CRC molecular subtypes (Figure 2B,C), we assessed whether snapshot measurement of individual tumour stem cell index was associated with clinical outcome. We did not observe any significant associations between quintiles of tumor stem cell index and survival in three CRC datasets after a Cox proportional hazards regression, indicating that snapshot measurement of stem cell index is not informative as a reliable, standalone prognostic molecular signature. However, in mouse and organoid models, temporally-spaced assessment of the stem cell index did detect dynamic and adaptive shifts in the tumor molecular phenotype in response to acute selective pressures. To undertake comparable assessment of molecular phenotypic shifts in human colorectal cancer, we turned to the Fluoropyrimidine, Oxaliplatin & Targeted Receptor pre-Operative Therapy for colon cancer (FOxTROT) trial data set (track A). This unique cohort of patients were randomised to 6 weeks neoadjuvant oxaliplatin and 5-FU treatment between diagnostic biopsy and surgical resection, enabling temporally-spaced transcriptional analysis of the tumor in response to a therapeutic selective pressure (Figure 6B) (Seymour *et al*., 2019). A spectrum of response to therapy could be identified, ranging between negligible and dynamic shifts in the stem cell index, and patients were grouped into “static” or “plastic” groups respectively (Figure 6C). Patients with “plastic” stem cell phenotype had no correlative change in transcriptome-based cell proliferation score (Figure 6D), but were significantly less likely to have a histologically detectable response to chemotherapy, indicating an association between the tumor capacity for adaptive change and clinical response to treatment (Figure 6E).

**Figure 6.**
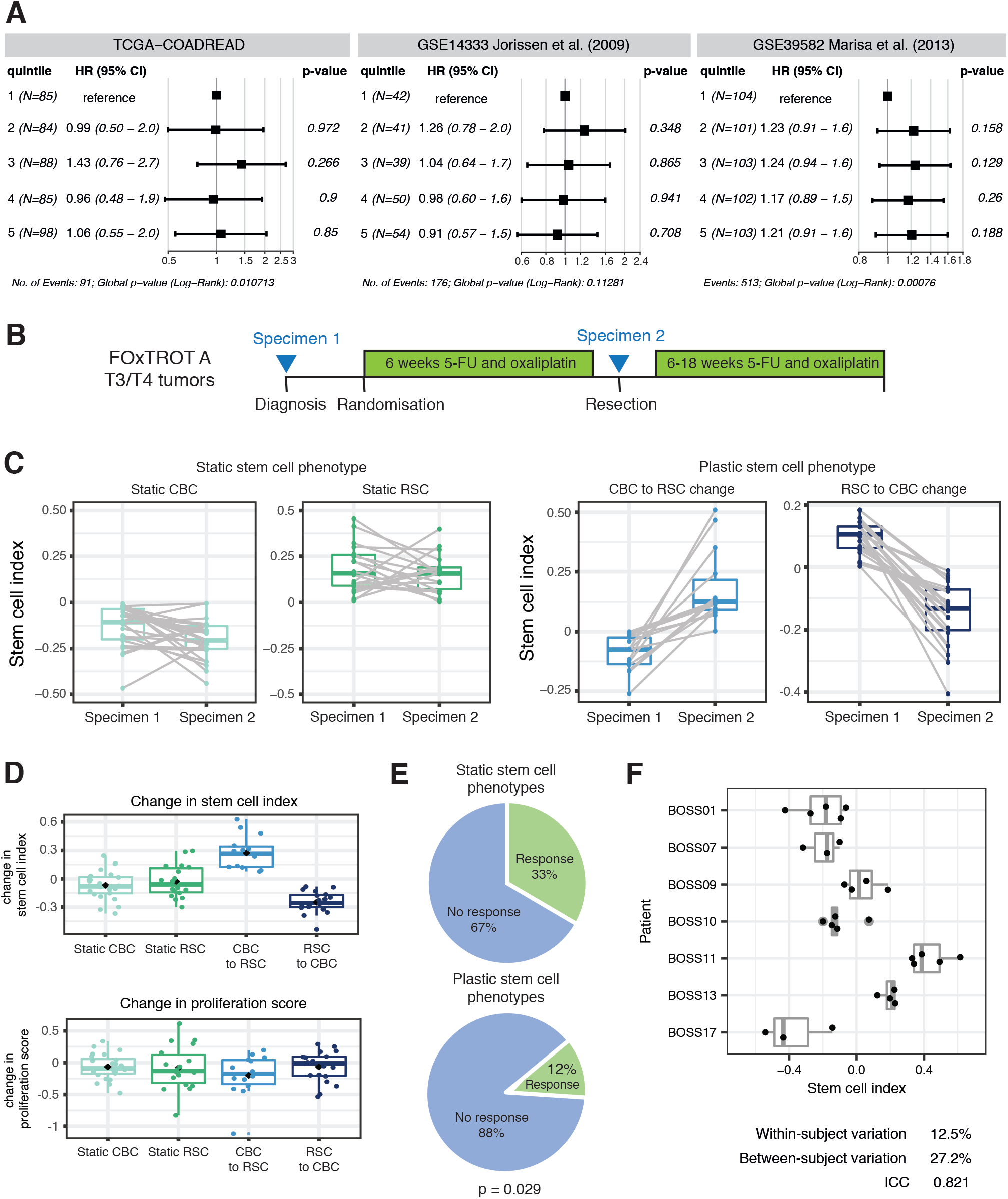
Human translational implications. **A**. Forest plots of progression free survival (PFS) HR (TCGA-COADREAD) and disease-free survival (DFS) HRs (Jorissen *et al*., 2009 and Marisa *et al*., 2013) for quintiles of tumor stem cell index. Data are presented as HR with error bars indicating the 95% CI. *P* values from a Cox proportional hazards regression are shown. **B**. FOxTROT (track A) trial schedule showing specimen acquisition before (specimen 1) and after 6 weeks (specimen 2) of 5-FU and oxaliplatin chemotherapy. **C**. Ladder plots showing GSVA (RSC-CBC signature) analysis of human tumor samples before and after chemotherapy, with patients grouped as “static” or “plastic” dependent on magnitude of signature change following therapy. **D**. No change in cell proliferation score in groups of tumours segregated by post treatment shift in stem cell index. **E**. Proportion of patients with a documented response to chemotherapy in the FOxTROT trial when grouped by static or plastic stem cell response to treatment. Statistical analysis, Fisher’s exact test, p value as stated. **F**. Low within-subject variation of the stem cell index from random non-adjacent biopsies from the BOSS trial.

We considered intratumour heterogeneity with resultant variability between biopsy and resection specimens could be a key potential confounder when interpreting consecutive samples in FOxTROT. It has been noted that sample heterogeneity contributes to highly variable consensus molecular subtype assignment in multiregional biopsy samples (Ubink *et al*., 2017), and this has been attributed to different proportions of stromal and immune cells in different biopsies (Alderdice *et al*., 2018). To assess this, we analysed stem cell index in the transcriptome data derived from non-adjacent, multiregional biopsies in the Biopsies of Surgical Specimens (BOSS) study (Ubink *et al*., 2017) and found a relatively high intra-class correlation coefficient (ICC = 0.821, Figure 6F). The within-subject coefficient of variation (variation between random biopsies from the same tumour, 12.5%) was lower than the between-subject coefficient of variation (27.2%), indicating that measurement of the stem cell index can transcend sample ‘contamination’ with non-epithelial tissue. Thus we concluded that measurement of the stem cell index from biopsy samples produces a reasonably robust representation of the stem cell phenotype of the whole tumour, which is reassuring for the conclusions drawn from temporally assessed samples in FOxTROT.

Together these data indicate that CBC and RSC marker-expressing cell populations can co-exist in intestinal neoplastic lesions, and that plastic cells can adaptively shift between these stem cell phenotypes in response to transgenic ablative or therapeutic selective pressures. Importantly, it is the capacity of the tumor to adapt, assessed through a responsive shift in the stem cell phenotype, rather than a snapshot assessment of the pre-treatment phenotypic equilibrium set point that is associated with chemoresponsiveness.

## Discussion

In the injured intestine, cell de-differentiation is an evolved, physiological response to enable rapid epithelial regeneration through temporary disruption of morphogen signalling and relaxation of stringent homeostatic controls over cell fate. Despite the neoplastic co-option and corruption of the same conserved signalling pathways, the contribution of analogous stem cell plasticity in colorectal tumors has not been fully established. This has clinical relevance, as cellular plasticity and resultant phenotypic heterogeneity drives therapy evasion, which has predominantly been noted in tumours where there is a definable morphological phenotypic switch — such as neuroendocrine differentiation in castration resistant prostate cancer and small cell conversion in treated nonsmall cell lung cancer (Boumahdi and de Sauvage, 2020). Assessment of this phenomenon in colorectal cancer has been hampered by the lack of a defined, measurable and pathologically-relevant phenotypic state that can temporally track lineage fate.

Here, we have used a combined molecular and morphological approach to assess stem cell phenotypic heterogeneity across a wide range of mouse and human intestinal tumors, and have developed an applicable and accessible tool to measure it – the stem cell index. This tool can be deployed on bulk tumor samples and is capable of detecting variable enrichment and admixture of CBC and RSC transcriptional signatures across lesions. This indicates that different populations of stem cells can co-exist in both benign and malignant neoplastic settings. Although not all cells actively expressing established stem cell markers necessarily serve as functioning stem cells (Barriga *et al*., 2017; Kozar *et al*., 2013), we were able to demonstrate comparable *ex vivo* stem cell potential in disaggregated primary tumor cell populations following FACS enrichment for CBC and RSC cell markers. Directional skewing of phenotype towards CBC or RSC predominance correlates with specific driver mutation(s) and/or disruption of microenvironmental signalling, with remarkable conservation of identified key pathways across mouse and human tumors. Using mouse models to control for epithelial mutation, we have generated data to support a coevolutionary, crosstalk model of solid tumor carcinogenesis, where epithelial accumulation of somatic mutation results in remodelling of the tumor context to establish genotype distinct niches, comprising variable immune, stromal and matrix components. Key morphogen pathways associated with these distinct microenvironmental cell compartments then signal back to the epithelium to induce cell plasticity and adaptively regulate stem cell phenotype. Although we show correlation of different stem cell phenotypes with some key signalling pathways, and note the overlap with the mechanisms that regulate wnt independence in murine organoids (Han *et al*., 2020), it is likely that multiple convergent and redundant mechanisms are involved in this intercompartmental crosstalk. Further detailed work, using advanced biological models, is needed to identify, map and therapeutically manipulate the combinatory networks of cell-intrinsic and extrinsic signalling disruption that induce, regulate and constrain stem cell plasticity in different tumor settings.

Consistent with a role for bidirectional epithelialmicroenvironmental crosstalk in regulating stem cell phenotype, we use the stem cell index to demonstrate rapid and measurable adaptive change in response to media manipulation of microenvironmental signalling *in vitro*, or an ablative direct selective pressure to crypt-base columnar cells *in vivo*. Loss of one stem cell population *in vivo* provokes cell plasticity and an acute adaptive switch in phenotype, with subsequent rapid recovery and restitution of the ablated CBC stem cell phenotype, restoring an equilibrated heterogeneic stem cell population. This is consistent with data from recently published models of secondary tumors. Circulating metastatic cells were noted to be predominantly *Lgr5*-ve, however, cell plasticity with *Lgr5*+ve stem cell reconstitution in the liver was required for secondary outgrowth (Fumagalli *et al*., 2020). Furthermore, constitutive YAP activation, with resultant continuous drive towards a RSC phenotype, prevented re-establishment of a stem cell equilibrium and abrogated orthotopic xenograft tumor or metastatic outgrowth (Cheung *et al*., 2020; Heinz *et al*., 2022). Together, this indicates the importance of a heterogeneous and dynamic stem cell population to enable adaptive response to selective pressures and to facilitate lesion outgrowth.

Here, in primary tumors, we demonstrate restoration of the stem cell equilibrium within 5 days of selective ablation, through rapid induction of cell plasticity and adaptive phenotypic shift. Deployment of the stem cell index was able to detect this profound change in the stem cell phenotype, but interestingly, mean lesion size in the animals was unaffected, and cell proliferation continued unchecked. A comparable dynamic change was detected in a proportion of patients in the valuable FOxTROT cohort, where serial sampling permits assessment of temporal change under the influence of a chemotherapeutic selective pressure. In this unique trial cohort, the use of temporally spaced stem cell index assessments acts as a measure of tumor adaption, and demonstrates that the capacity for dynamic change in stem cell marker expression was associated with reduced histological response to therapy. These combined preclinical and human data demonstrate that induced shifts in the molecular phenotype may occur independently of the response seen using conventional assessments of clinical disease activity, such as tumor size or cell proliferation. Stem cell molecular phenotype may be an informative metric that is currently going unmeasured when considering patient response to therapy. The capacity to temporally track cell fate and potentially titrate treatments, according to detectable changes in a disease relevant molecular phenotype could become a powerful tool in patients undergoing neoadjuvant therapies, such as chemoradiotherapy in rectal cancer.

Using the data presented here we propose a modified fitness landscape model of stem cell phenotype in colorectal cancer, where *Lgr5*+ve CBC and *Lgr5*-ve RSC represent distinct, but interlinked and equilibrated stem cell population peaks situated along a phenotypic axis. Epithelial cells shift their stem cell phenotype along this axis through the combination of acquired epithelial mutation and the influence of microenvironmental signalling (Figure 7A), which arises from surrounding niches comprising of variably remodelled immune, stromal and matrix tissue compartments. Establishment of a tumor-specific stem cell population equilibrium depends on the balance of cell-intrinsic and extrinsic signalling. Application of a selective pressure to a stem cell phenotype peak alters microenvironmental morphogenic signalling resulting in a new fitness landscape, and temporarily shifts the stem cell population distribution towards an alternative phenotype. This cell plasticity allows reconstitution of the lost stem cell population, facilitating a recovery of the stem cell distribution to a new, post-treatment equilibrium (Figure 7B). Critically, it seems to be the capacity of the tumor to enable this adaptive phenotypic shift and recover an equilibrated heterogeneic stem cell population that is associated with chemoresistance, rather than the pretreatment position of the phenotypic equilibrium set point. This is consistent with recently published work that shows that constitutive YAP activation drives a polarised regenerative stem cell phenotype and prevents restoration of the dynamic and heterogeneous stem cell population required for lesion outgrowth (Cheung *et al*., 2020; Heinz *et al*., 2022). In our model, this can be represented by skewing of the stem cell phenotype onto a single population peak which constitutes an evolutionary dead end.

**Figure 7.**
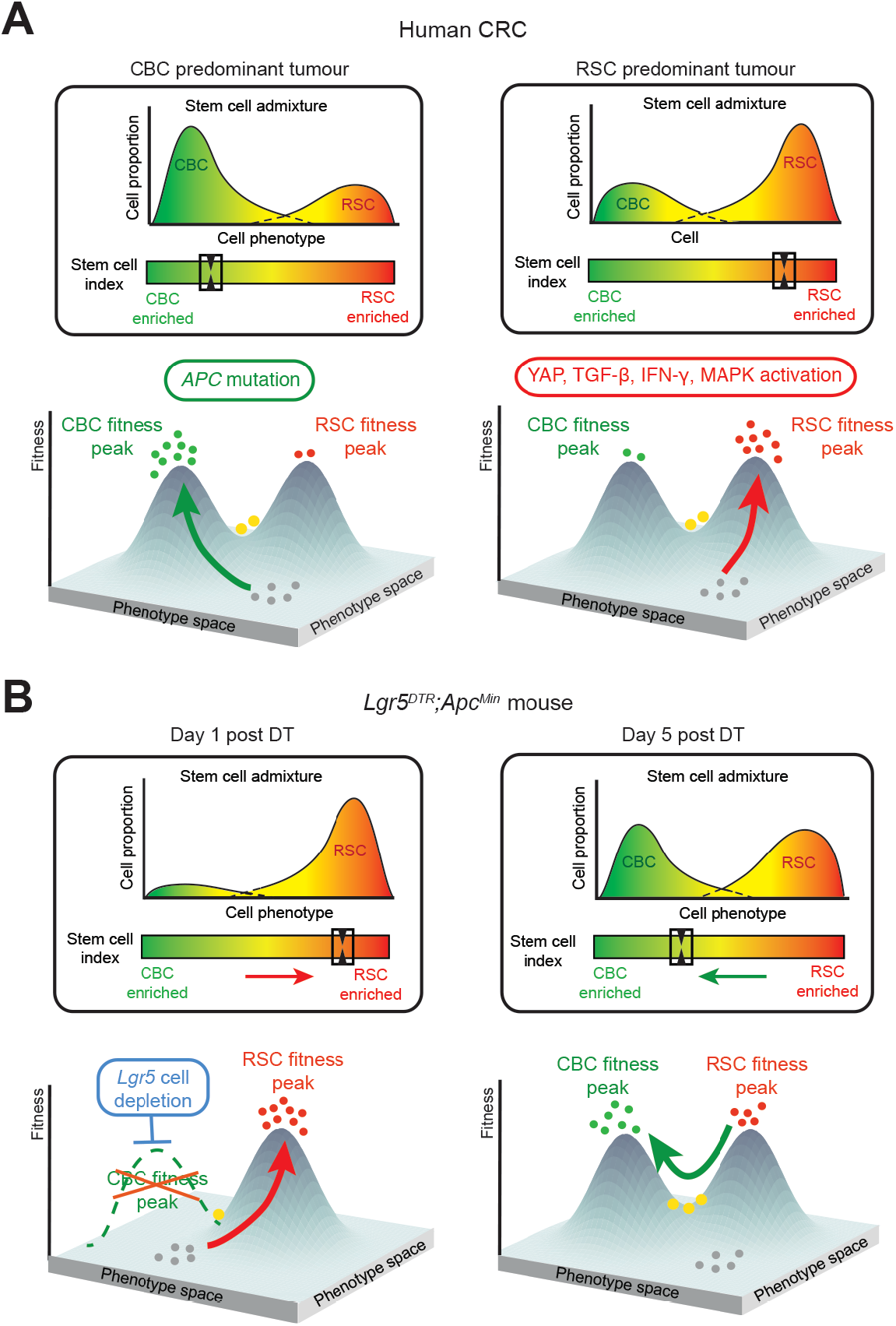
Fitness landscape model. Tumor stem cell phenotype can be represented as a fitness landscape model, where *Lgr5*+ve CBC and *Lgr5*-ve RSC represent distinct, but interlinked fitness peaks situated along a phenotypic axis. **A**. Epithelial cells ‘climb’ these fitness peaks (arrows) through the combination of acquired epithelial mutations and the influence of microenvironmental signalling placing them at distinct points within the fitness landscape. Bulk transcriptome data can be used to calculate the stem cell index which reflects stem cell population admixture and can be used as a measure of individual tumour position within this phenotypic axis (boxes) **B**. Application of a selective pressure to a fitness peak (e.g. *Lgr5+* CBC ablation), alters the morphogenic signalling landscape and shifts the stem cell equilibrium towards an alternative phenotype at day 1. Rapid regeneration of the lost CBC stem cell population restores the stem cell equilibrium after 5 days. These dynamic shifts can be measured by change in the stem cell index (boxes). Key: green dots CBC cells, red dots – RSC cells, yellow dots – both marker expressing cells, grey dots – no stem cell marker expressing cells.

We have developed a simple and applicable tool to assess stem cell admixture and detect responsive shifts in the stem cell molecular phenotype. We believe that this can be used to biologically steer neoadjuvant therapies to inform the scheduling of our current therapeutic armamentarium. Concomitant development of new treatments is needed, both to target stem cell populations directly, and to close off the emergent adaptive signalling pathways that enable stem cell plasticity in order to abrogate the evolution of therapy resistance.

### Limitations of the study

We have not undertaken whole exome somatic mutation screening of all of the lesions from the autochthonous animal models, thus it is conceivable that some of the stem cell index intra-genotype variation seen (particularly in the KPN group), is the consequence of undetermined additional somatic mutation. Furthermore, the complex multi-allele mouse models used here currently result in simultaneous genetic recombination. Thus, changes seen in the genotypespecific tumour landscape remodelling are the result of concomitant, combined driver gene mutation rather than the cumulative, stepwise progression seen in human disease.

## Supporting information

Supplementary Table 1 - RSC gene list

Supplementary Table 2 - CBC gene list

## Acknowledgements

The authors thank Genentech (California, USA) for the supply of the *Lgr5*^*DTR*^ mouse.

This paper was typeset with the bioRxiv word template by @Chrelli: www.github.com/chrelli/bioRxiv-word-template

## Author contributions

EGV, NN and SJL conceived and designed the project. Funding obtained by SJL, OS. Experiments were conducted by EGV, NN, HB-D, EM, TRML, DF, AKN, SO, AM, KS, AC, VW, DH. Bioinformatic analysis carried out by GV, KG, PW, PT, ED, NN, EGV. Pathology support, image analysis and intellectual input from VHK, AE. Tissue and data provision JEE, ST, TM, DM, DC, LT. Conceptual input and data interpretation OS, PD, ST. Manuscript written by EGV, NN, GV and SJL.

## Disclaimer

The views expressed are those of the author/s and not necessarily those of the NHS, the NIHR or the Department of Health.

## Materials and Methods

### Resource availability

Further information and requests for resources and reagents should be directed to and will be fulfilled by the lead contact, Simon Leedham.

### Materials Availability

Mouse lines and organoids generated in this study are available with an MTA.

### Data and Code Availability

The RNA-seq data generated for this paper have been deposited at the ArrayExpress database at EMBL-EBI (www.ebi.ac.uk/arrayexpress) under accession numbers E-MTAB-10470, E-MTAB-11769, and E-MTAB-11784, and are publicly available as of the date of publication.

A function to calculate the stem cell index has been made available in an R package *ISCindex* (https://github.com/gnvalbuena/ISCindex). All original code has been deposited at Zenodo (DOI: 10.5281/zenodo.6473396) and is publicly available as of the date of publication.

Any additional information required to reanalyze the data reported in this paper is available from the lead contact upon request.

### EXPERIMENTAL MODEL AND SUBJECT DETAILS

#### Animals

The study comprised the use of two mouse cohorts: an internal mouse i.e. Oxford cohort and the ACRCelerate cohort. All procedures were carried out in accordance to Home Office UK regulations and the Animals (Scientific Procedures) Act 1986. All mice are housed in individually ventilated cages at the animal unit either at Functional Genetics Facility (Wellcome Centre for Human Genetics, University of Oxford) or The Beatson Institute (Glasgow). All mice were housed in a specific-pathogen-free (SPF) facility, with unrestricted access to food and water, and were not involved in any previous procedures. All strains used in this study were maintained on C57BL/6J background for ≥6 generations. All procedures were carried out on mice of at least 6 weeks of age, both male and female.

#### Human subjects

Samples were collected from 4 cohorts: S:CORT cancer cohort, Oxford BRC cancer cohort, pre-cancer polyps cohort and colitis cohort. All human samples were obtained following ethical approval and individual informed consent (Ethics No 16/NI/0030 and 15/EE/0241 for S:CORT cohorts, REC: 17/NW/0252 BRC cancer cohort, and Oxford GI biobank ethics 18/SL/JE/Early-detection and OCHRe:18/A11 for the colitis cohort). All samples were subject to expert histopathological review. Data from the FOXTROT trial, developed by the NCRI Colorectal Cancer Clinical Studies Group (NCT00647530) has been used in this study.

### METHODS DETAILS

#### Treatment of Animals

The mouse alleles used in this study can be used in the Key Resources Table. Phenotype induction was obtained by intraperitoneal tamoxifen (Merck) injection in the inducible models. *Lgr5*+ cells in the mice were ablated with a single intraperitoneal dose of Diphtheria Toxin (DT) (Cat # 322326-1mg, Merck) in saline (50 µg kg^−1^).

#### FFPE Processing

Gut preparations were washed in PBS, fixed overnight in 10% neutral buffered formalin and then transferred to 70% ethanol prior to processing for embedding. Formalin**-**fixed gut sections were rolled into Swiss Rolls, pinned and placed in a histology cassette. Specimens were processed using a Histomaster machine (Bavimed). Processed samples were embedded in paraffin wax using a paraffin embedding station (EG1150H, Leica).

### Nucleic acid extraction

For the Oxford mouse cohort, RNA was extracted using the RNeasy Micro Kit from Qiagen (74004) and DNase treatment was performed using the DNA-free kit from Life Technologies (AM1906).

For the ACRCelerate mouse cohort, RNA extraction was performed using a Qiagen RNeasy kit according to the manufacturer’s protocols.

For FFPE samples from human cohorts, 2-10 5 micron sections were extracted using the High Pure FFPE RNA Isolation kit (Roche Life Sciences, Penzberg, Germany) under RNase free conditions following the manufacturer’s protocol. RNA quantity and quality were assessed using the RNA Qubit Assays (High sensitivity and Broad range, Thermofisher) and by Nanodrop, respectively.

### RNA sequencing

Transcriptomic profiling was carried out by 3’RNAseq for the Oxford mouse cohort, pre-cancer polyps and the colitis cohorts. Libraries were sequenced on an Illumina HiSeq4000 instrument (Illumina). 3’RNAseq libraries were prepared using the QuantSeq 3’ mRNA-Seq Library Prep Kit FWD for Illumina (Lexogen, Austria). The manufacturer’s protocol for FFPE or fresh frozen tissue was used based on the source tissue material. For murine sample libraries, RNA input varying from 20-1000ng and 14-20 PCR cycles were used. For human sample libraries, RNA input varying from 53-10,000ng and 16-21 PCR cycles were used. Library quality and quantity were assessed using the HS 1000 DNA TapeStation (Agilent) and DNA High Sensitivity Qubit Assay (ThermoFisher) respectively, before pooling and sequencing on an Illumina HiSeq4000 instrument (Illumina).

For the ACRCelerate cohort, libraries were prepared using a TruSeq RNA sample prep kit v2 (Illumina) and sequenced on an Illumina NextSeq using the High Output 75 cycles kit (2x36 cycles, paired end reads, single index). For the ACRCelerate cohort, raw sequence quality was assessed using FastQC version 0.11.8, then sequences were trimmed to remove adaptor sequences and low-quality base calls, defined as those with a Phred score of less than 20, using Trim Galore version 0.6.4. Trimmed sequences were aligned to mouse genome build GRCm38.98 using HISAT2 version 2.1.0 and raw counts per gene were determined using FeatureCounts version 1.6.4. Counts were then normalised via quantile normalisation in R.

### Microarray transcriptome profiling

The S:CORT cancer cohort transcriptomes were analysed by microarray. Extracted RNA was first amplified using the NuGen Ovation FFPE Amplification System v3 (NuGen San Carlos, California, USA). The amplified product was hybridised to the Almac Diagnostics XCEL array (Almac, Craigavon, UK), a cDNA microarray-based technology optimised for archival FFPE tissue, and analysed using the Affymetrix Genechip 3000 7G scanner (Affymetrix, Santa Clara, California, USA) as previously described (12). Microarray data were quality checked then pre-processed where raw CEL files underwent the Robust Multiarray Average (RMA) normalisation for the Almac Diagnostic XCEL array with the affy package (v1.56.0) (17).

### *In situ* hybridisation

For *in-situ* hybridisation (ISH) of both human and mouse FFPE samples, 4 µm formalin-fixed, paraffin-embedded tissue sections were used. The sections were baked at 60°C for 1 hour before dewaxing in xylene and ethanol. Fluorescent *ISH* was then performed using the RNAscope Fluorescent Multiplex Reagent Kit (Bio-techne) in accordance with the supplier’s guidelines. Probes were purchased from Bio-techne: Mm-Anxa1 (509291), HsANXA1 (465411), Mm-Ly6a-C2 (427571-C2), Hs-PLAUR (542701), MmClu-C3 (427891-C3), Hs-CLU (606241), Mm-Lgr5-C2 (312171-C2), Hs-LGR5-C2 (311021-C2).

### Immunohistochemistry

Sections were de-paraffinized in xylene and rehydrated through graded alcohols to water. Antigen retrieval was done by pressure cooking in 10mmol/L citrate buffer (pH 6.0) for 5 minutes. Endogenous peroxidase activity was blocked by incubating in 3% hydrogen peroxidase (in methanol) for 20 minutes. Next, sections were blocked with 1.5% serum for 30 min, after which they were incubated with primary antibodies for 2 h. Antibodies against the following proteins were used: KI67 (Cell Signaling Technology, CS12202S, 1/500), Caspase 3 (R&D Systems AF835, 1:800). The sections were then incubated with appropriate secondary antibodies for 30 minutes at room temperature. For chromogenic visualization, sections were incubated with ABC (Vector labs) for 30 min and stained using DAB solution (VectorLabs), after which they were counterstained with hematoxylin, dehydrated and mounted.

### Multiplex immunofluorescence

Multiplex immunofluorescence (MPIF) staining was performed on formalin-fixed paraffin embedded (FFPE) sections of thickness 4-µm using the OPAL protocol (Akoya Biosciences, Marlborough, MA) on the Leica BOND RXm autostainer (Leica Microsystems, Wetzlar, Germany). Six consecutive staining cycles were performed using the following primary antibody-Opal fluorophore pairs:

#### Immune panel

(1) Ly6G (1:300, 551459; BD Pharmingen)–Opal 540; (2) CD4 (1:500, ab183685; Abcam)–Opal 520; (3) CD8 (1:800, 98941; Cell Signalling)–Opal 570; (4) CD68 (1:1200, ab125212; Abcam,)–Opal 620; (5) FoxP3 (1:400, 126553; Cell Signalling)–Opal 650; and (6) E-cadherin (1:500, 3195; Cell Signalling)–Opal 690.

#### Stroma panel

(1) Gremlin 1 (1:750, AF956; R&D)–Opal 540; (2) CD34 (1:3000, ab81289; Abcam)–Opal 520; (3) CD146 (1:500, ab75769; Abcam)– Opal 570; (4) SMA (1:1000, ab5694; Abcam)–Opal 620; (5) Periostin (1:1000, ab227049; Abcam)–Opal 690; and (6) E-cadherin (1:500, 3195; Cell Signalling)–Opal 650.

#### Matrix panel

(1) Laminin (1:400, ab11575; Abcam)-Opal 540; (2) Tenascin-C(1:600, ab108930; Abcam)-Opal 520; (3) Fibronectin (1:1000, F3648; Sigma-Aldrich)-Opal 570; (4) Osteopontin (1:750, ab218237;Abcam)-Opal 620; MMP3 (1:100,ab52915; Abcam)-Opal 650; (5) Collagen I (1:400, 72026; Cell Signalling)-Opal 690.

Tissue sections were incubated for 1 hour in primary antibodies and detected using the BOND Polymer Refine Detection System (DS9800; Leica Biosystems, Buffalo Grove, IL) in accordance with the manufacturer’s instructions, substituting DAB for the Opal fluorophores, with a 10-minute incubation time and withholding the hematoxylin step. Antigen retrieval at 100ºC for 20 minutes, in accordance with standard Leica protocol, with Epitope Retrieval (ER) Solution 1 or 2 was performed prior to each primary antibody being applied. Sections were then incubated for 10 minutes with spectral DAPI (FP1490, Akoya Biosciences) and the slides mounted with VECTASHIELD Vibrance Antifade Mounting Medium (H-1700-10; Vector Laboratories). Whole-slide scans and multispectral images (MSI) were obtained on the Akoya Biosciences Vectra Polaris™. Batch analysis of the MSIs from each case was performed with the inForm 2.4.8 software provided. Finally, batched analysed MSIs were fused in HALO® (Indica Labs) to produce a spectrally unmixed reconstructed whole-tissue image. Cell density analysis was subsequently performed for each cell phenotype across the three MPIF panels using HALO^®^

### Tumour organoid generation and maintenance

Organoid growth media was made from advanced DMEM/F12 supplemented with penicillin (100 U/ml) and streptomycin (100 µg/ml) (ThermoFisher Scientific, 15140122), 2 mM L-glutamine (ThermoFisher Scientific, 25030081) 10 mM HEPES (ThermoFisher Scientific, 15630080), N2supplement (ThermoFisher Scientific, 17502001), B27-supplement (ThermoFisher Scientific, 17504044), recombinant human EGF 50 ng/ml (Peprotech, AF-100-15) and recombinant murine Noggin 100 ng/ml (Peprotech, 250-38).

Tumour fragments from mice were taken in ice cold PBS at time of dissection. Fragments were then cut into 2-5mm pieces and washed in ice cold PBS three times. These were then incubated in 5ml of 10x Trypsin and 200U recombinant DNase at 37°C for 30 minutes after being shaken vigorously. Next, 20ml of ADF was added and fragments were shaken vigorously again. Samples were spun and supernatant was aspirated. The pellet was resuspended in 10ml of ADF and passed through a 70um strainer. Tube and strainer were then rinsed with another 5ml of ADF. The filtered suspension was pelleted and supernatant aspirated. This was suspended in Matrigel and plated depending on pellet size. The plate was inverted and left to set at 37°C for 10 minutes before adding standard growth media. Organoids were grown at 37°C 5% CO2 21% O2.

Organoids were passaged every 2-3 days using mechanical dissociation and split 1:3 or 1:2 ratio depending on organoid density. Organoid lines were frozen in Gibco cell culture freezing media and revived before use for experiments. Routine mycoplasma testing was done before transplantation experiments. For RNA seq, organoids were washed in PBS, pelleted and snap frozen on dry ice.

### Organoid treatment with Interferon-gamma & TGF-β

Wild-type mouse colonic organoids were plated onto 24-well plates in Matrigel and allowed to culture for 5 days before treatment with recombinant murine INF-y (PeproTech, UK) at concentrations of 0.2, 1, and 5 ng/mL. 24 h after treatment RNA was extracted and cDNA synthesised, and the target genes Lgr5 and Ly6a, GAPDH were measured by qRT-PCR.

Mouse intestinal organoids (wild-type, AKPT, and KPN) were plated onto 24-well plates in Matrigel and cultured for 3 days before treatment with either recombinant murine INF-y (PeproTech, UK) at 1 ng/mL or recombinant murine TGFβ 1 (R&D, UK) at 5 ng/mL. RNA was extracted and processed for RNA sequencing 24 h after treatment.

### FACS sorting of IFN-γ treated Lgr5-GFP organoids

For FACS analysis of IFN-γ-treated organoids, adult *Lgr5;EGFP;CreERT2* organoids were grown in matrigel as previously described (Sato, Nature 2009). Interferon gamma (R&D rmIFN-gamma #485-MI, 5ng/ml) was added on day 5 after passaging. After 24hrs, organoids were dissociated into single cells by incubation with 1ml TrypLE Express (Gibco) for 15 mins at 37C with mechanical dissociation after 10 mins. Cells were then incubated with anti-mouse Ly6a-APC (1/1000, eBioscience #17-5981-81) for 30 mins on ice. Hoechst 33342 was added to exclude live from dead cells and an Isotype control (APC) was used for gating.

### Stem cell sorting and single cell clonogenicity analysis

Stem cell populations were sorted from KPN mice. Mice were sacrificed upon development of intestinal phenotype. The intestinal tissue was removed, and subjected to manual dissociation followed by enzymatic dissociation with 5mL 10x Trypsin (5mg/ml, Gibco), 1x DNase buffer (500ul) and 200U recombinant DNase I (20ul, Roche, 04716728001) at 37 C in a shaker set at 100 RPM. The cell suspension was then The cell suspension was first incubated in Mouse Fc Block purified anti-mouse CD16/CD32 mAb, then stained for the following markers: CD31, CD45, EpCAM, Sca1 and Ephb2. The stained cells were sorted for the following populations on a BD FACSAria Fusion Flow Cytometer: (1) Sca1+Ephb2+, (2) Sca1+Ephb2-, (3) Ephb2+Sca1-, and (4) Sca1-Ephb2-. The gating strategy used is shown in Figure S2. The cells were collected in media on 1mL Eppendorf tubes before being grown in matrigel supplemented with Jagged-1 (Anaspec, 1 µM) and cultured in advanced DMEM/F12 (Thermo Fisher Scientific) supplemented with GlutaMAX (Thermo Fisher Scientific, 1%) and Penicillin/Streptomycin (Thermo Fisher Scientific, 0.5%) in the presence of human EGF (Peprotech; 50ng/mL), murine Noggin (Peprotech; 100ng/mL), human R-spondin1 (R&D; 500ng/mL), murine Wnt3a (Cell guidance systems; 100ng/ml), Chir99021 (Stemgent; 3 microM), Prostaglandin E2 (PGE2; Sigma; 2.5microM), Nicotinamide (Sigma; 10 mM), N-2 supplement (Thermo Fisher Scientific; 1%) and B-27 supplement (Thermo Fisher Scientific; 2%). In the first 3 days, cells were provided with Y-27632 as additional supplement. Cells were imaged at Day 7 from plating and number of organoids manually counted.

## QUANTIFICATION AND STATISTICAL ANALYSIS

### Calculation of the Stem Cell Index

The RSC gene signature was derived from the repair signature in Yui *et al*. (2018) and the spheroid upregulated gene list from Mustata *et al*. (2013). Cell type-associated expression was characterised using hierarchical clustering of signature genes in published single cell RNAseq data from colorectal tumors and normal colon tissue (Lee *et al*., 2020), and genes from clusters that represented epithelial-expressed genes with high stromal expression, stromal-expressed genes with no distinctive expression in other cell types, as well as genes with no clear cell-type specific expression were excluded from the signature. The resulting gene signature of 265 genes is provided in Supplementary Table 1.

The CBC signature was taken from the *Lgr5* intestinal stem cell signature in Muñoz *et al*. (2012).

Single sample enrichment scores for the RSC and CBC signatures were calculated using Gene Set Variation Analysis (GSVA, Hänzelmann *et al*., 2013) from TMM-normalised CPM values in RNAseq data and from mean per gene values in microarray data. The Stem Cell Index was calculated by subtracting the CBC score from the RSC score.

### Quantification of Casp3 and Ki67 positive cells

Casp3 and Ki67 positive cells were quantified using QuPath digital pathology software (v0.2.3, Bankhead *et al*., 2017), downloaded from https://QuPath.github.io/. Firstly, annotations of polyp areas were created for each tissue sample with areas of folded tissue excluded to eliminate false positive signals. Cells were identified within QuPath using a custom algorithm established via stain separation using colour reconstruction. Positive cell detection analysis was run to identify DAB positive cells for both Casp3 and Ki67 stained sections and results reported as percentage of positive cells. Each annotation was manually verified for correct signal identification.

### Gene set enrichment analysis

Gene Set Enrichment Analysis was performed using the *fgsea* package. Single sample enrichment scores were calculated using Gene Set Variation Analysis (GSVA, Hänzelmann *et al*., 2013).

The signatures used were previously published: Yap (Gregorieff *et al*., 2015; Yui *et al*., 2018), Wnt (Sansom *et al*., 2004; Van der Flier *et al*., 2007), Kras (Loboda *et al*., 2010), IFN-γ (Ayers *et al*., 2017), Fibroblast TGF-β response (Calon *et al*., 2012), and the MSigDB Hallmark gene set *(Liberzon et al*., 2015) and Reactome (Jassal *et al*., 2020). Mouse genes were mapped to the most confident human orthologs, based on the Ensembl annotation (March 2020).

### Single cell RNAseq analysis

Previously published human colon single cell RNA sequencing data was used (Lee *et al*., 2020, GEO: GSE132257 and GSE144735). The raw expression matrix from the *CellRanger* pipeline was normalised with *Seurat*. Cell type assignment was performed with *SingleR* (Aran *et al*., 2019). Epithelial cells were retained for downstream analysis and quality control filtering was applied for >1000 unique molecular identifier counts (UMI), <30% mitochondrial gene expression, minimum 2000, and maximum 6000 detected genes. Dimension reduction was performed with UMAP of highly variable genes, defined by variance modelling (function *modelGeneVar()*) at an FDR<5%, after excluding mitochondrial and ribosomal genes. We applied the mutual nearest-neighbour algorithm for dataset integration (Haghverdi *et al*., 2018).

Mouse genes were mapped to the most confident human orthologs, based on the Ensembl annotation (March 2020). The genes were further classified into epithelial, non-epithelial and non-specific, according to the specificity of their expression in the KUL3 colorectal single-cell dataset (Lee *et al*., 2020). Briefly, we aggregated raw counts from cells of similar type (epithelial, stroma, myeloid, lymphocytes, endothelial and other) into pseudobulk with the *muscat* package. If the expression of a gene was higher in a given cell type than in all other cell types by at least 1 raw count and that difference was always statistically significant at P<0.01, that gene was considered specific for that cell type. Thus, only genes of predominantly epithelial origin were kept from every signature. The mean expression value was used as the signature value for every cell.

The proportion of true LGR5 positive cells was expected to be close to 5%, according to previous studies (Cao *et al*., 2020). Therefore, the cutoff for LGR5 positivity was fixed at the 95th percentile of the signature values in the cell population (rounded to 0.45 logcounts). There was no *a priori* expectation for the proportion of true SCA1 positive cells, so on the assumption that the number of SCA1 positive stem cells would be comparable, the cutoff for SCA1 positivity was also fixed at the 95th percentile (rounded to 0.65 logcounts). Double-positive (“mixed”) cells were defined as having both SCA1 and LGR5 expression above the predefined cutoffs

### Mutational and Copy Number Alteration Analysis on TCGA colorectal cohort

Mutation Annotation Format files generated by SomaticSniper and mean segment copy number calls from the TCGA Colorectal adenocarcinoma project (TCGA-COADREAD) were downloaded from the GDC data portal. Mutations were filtered for pathogenicity using the *SIFT* and *Polyphen* predictors and subsequent analysis were conducted using R and results were plotted using the R function *barplot*. For copy number analysis, CNV values smaller than -0.3 were categorised as a loss (-1), whereas values above 0.3 were annotated as gains (+1). Gains and losses were plotted using the *plotFreq* function from the R package *copynumber*.

### Survival analyses

TCGA-COADREAD RNA sequencing and clinical data were downloaded from the Genomics Data Commons using TCGAbiolinks on R. Gene expression microarray and clinical data from GSE14333 (Jorissen *et al*., 2009) GSE39582 (Marisa *et al*., 2013) were downloaded from GEO. Patients with Stage 1-3/Dukes A-C cancers and with no missing data in any covariate were included in the analysis 440 in the TCGA-COADREAD data, 226 in the Jorissen *et al*. data, and 513 in the Marisa *et al*. data. The relationship between quintiles of stem cell index and survival (Progression-free survival/PFS in the TCGA-COADREAD data and Disease-free survival/DFS in the Jorissen *et al*. and Marisa *et al*. data) was examined using multivariate Cox regression analyses with TNM/Dukes stage, age, and gender as possible confounding factors. Survival analyses were performed in R using the survival package.

### Analysis of therapy response

Associations of stem cell phenotype to therapy response were investigated in data from the FOXTROT trial. The analysis was undertaken on 75 patients from Trial Arm A, who were given 6 weeks of pre-operative OxFP chemotherapy followed by surgery then 18 (or 6) weeks post-operative OxFP chemotherapy. For each of the patients, pre-treatment biopsies and a post-treatment resection sample were collected for genetic and transcriptomic profiling.

Response to treatment of patients was determined by histological assessment. Patients with no regression (no change) or mild regression (residual cancer cells are higher than fibrosis) were classified as non-responders, while those with marked regression (presence of fibrosis with rare residual cancer cells) or moderate regression (increased presence of residual cancer cells but still high fibrosis) were classified as responders.

The stem cell index was for each sample. Samples with a positive Index score were classified as RSC-positive. Samples with a negative Index score were classified as CBC-positive. Patients were discerned into: those with a static stem cell phenotype (where preand post-treatment samples were either both RSC-positive or both CBC-positive) and those with a plastic stem cell phenotype (where the tumour shifted either from being RSC-positive pre-treatment to CBC-positive post-treatment, or from CBC-positive pretreatment to RSC-positive post-treatment). Differences in treatment response between static and plastic stem cell phenotypes were evaluated using a Fisher’s exact test.

**Supplementary Figure 1.**
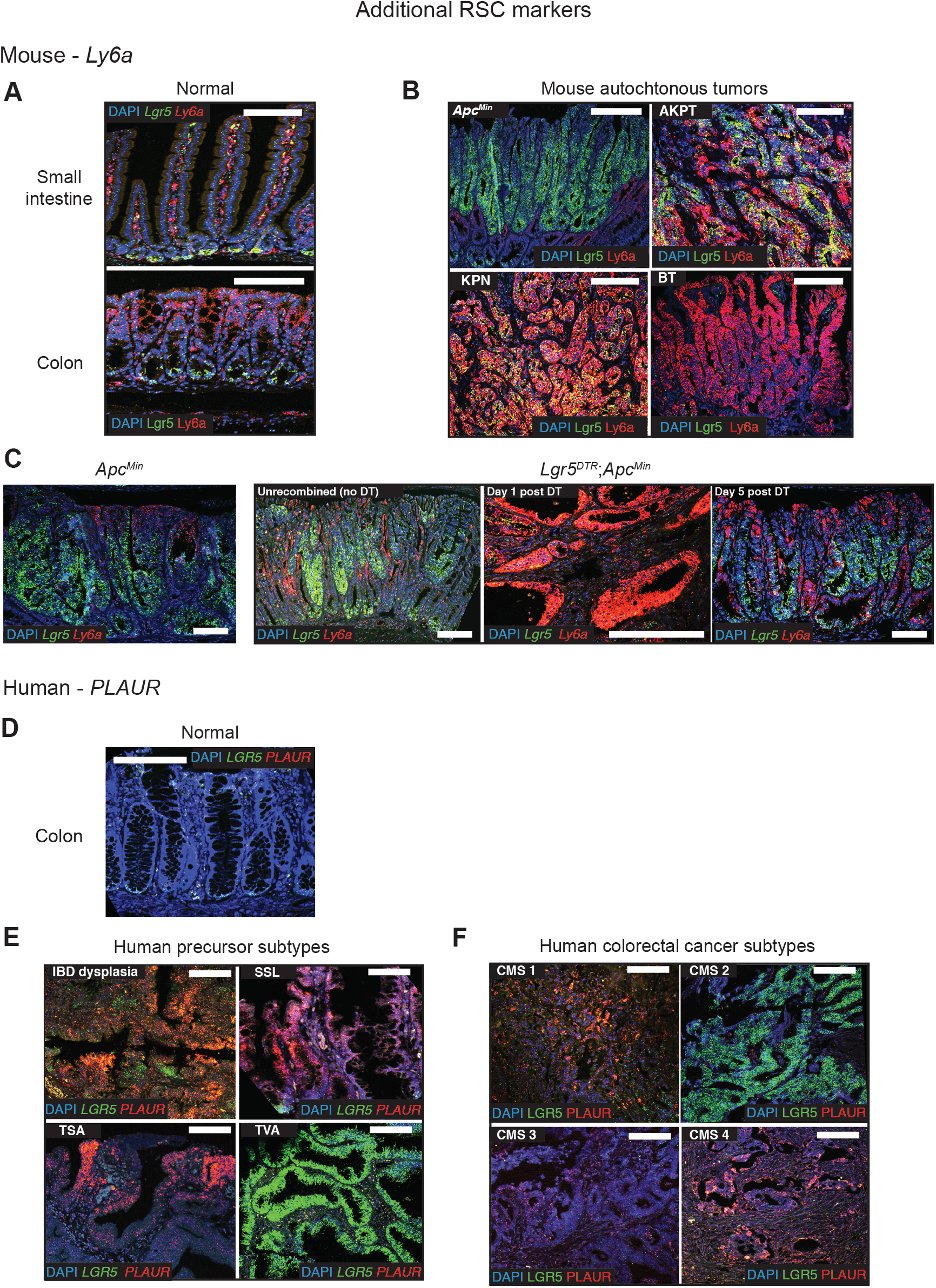
Additional RSC morphological markers. **A**. Dual color ISH for *Lgr5* (CBC marker - green) and *Ly6a* (RSC marker - red) expression in normal mouse small intestine and colon. **B**. Dual color ISH for *Lgr5* (CBC marker - green) and *Ly6a* (RSC marker - red) expression in representative genotype tumors across the CBC to RSC spectrum. **C**. Dual color ISH for *Lgr5* (CBC marker - green) and *Ly6a* (RSC marker - red) to show marker expression change before and after CBC cell ablation in *Lgr5*^*DTR*^*;Apc*^*Min*^ mice. Driver alleles initialisation - A is *Apc*^*fl/+*^, *Apc*^*Min*^ is *Apc*^*Min*^, B is *Braf*^*V600E*^, K is *Kras*^*G12D*^, P is *p53*^*fl/fl*^, T is *Tgfβr1*^*fl/fl*^, N is *Rosa26*^*N1icd/+*^ **D**. Dual color ISH for *LGR5* (CBC marker - green) and *PLAUR* (RSC marker - red) expression in normal human colon. **E-F**. Dual color ISH for *LGR5* (CBC marker - green) and *PLAUR* (RSC marker - red) expression in **E**. representative human precursor lesions and **F**. representative human colorectal cancers segregated by consensus molecular subtype.

**Supplementary Figure 2.**
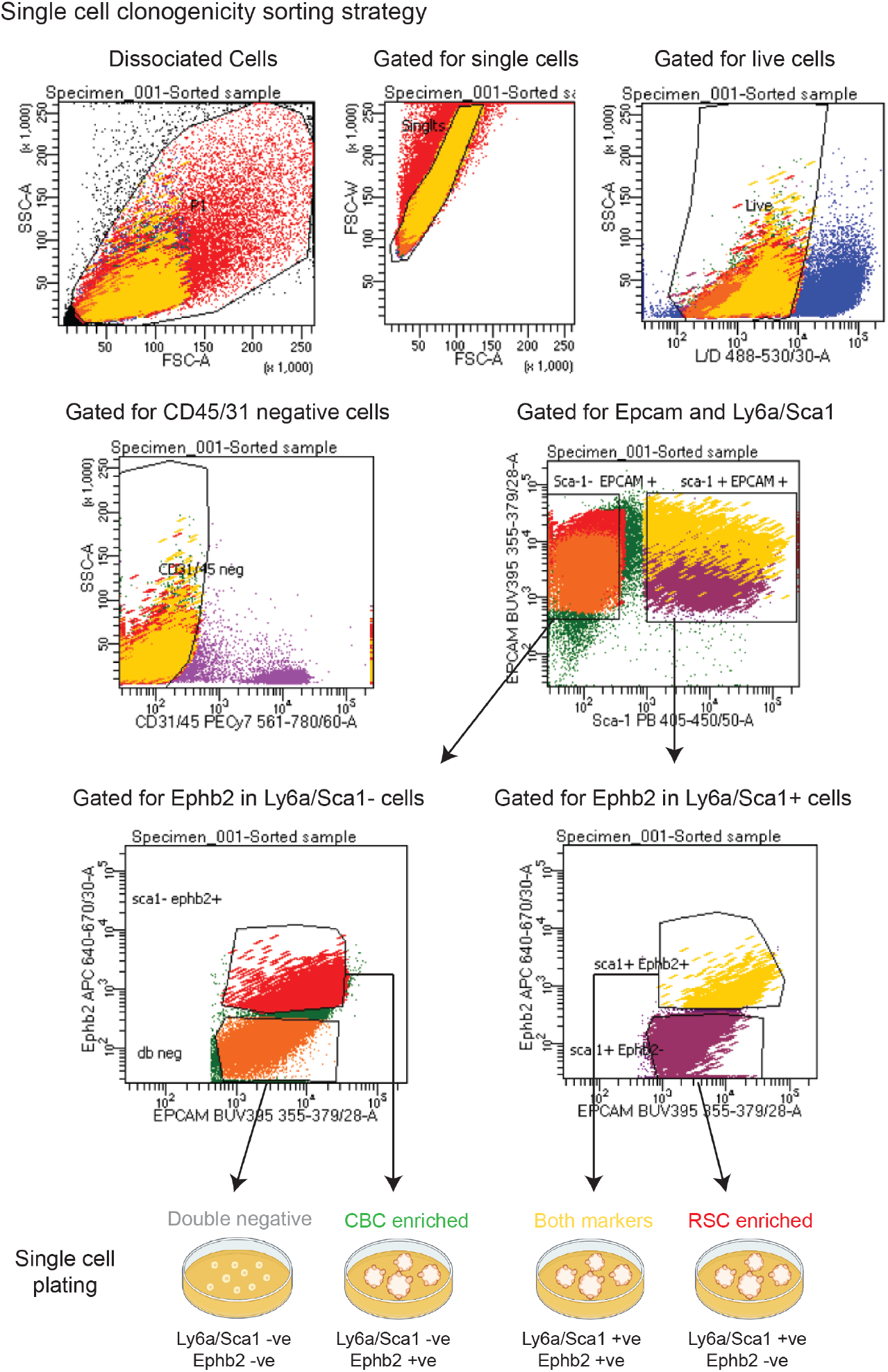
Single cell clonogenicity sorting strategy. KPN mouse primary tumours (n=5) were disaggregated and cells sorted to generate live epithelial cells in four distinct groups, CBC-enriched, RSC-enriched, expression of both CBC and RSC markers, and double negative cells. Sorted single cells were plated in Matrigel and organoid cloning efficiency was assessed 7 days later.

**Supplementary Figure 3.**
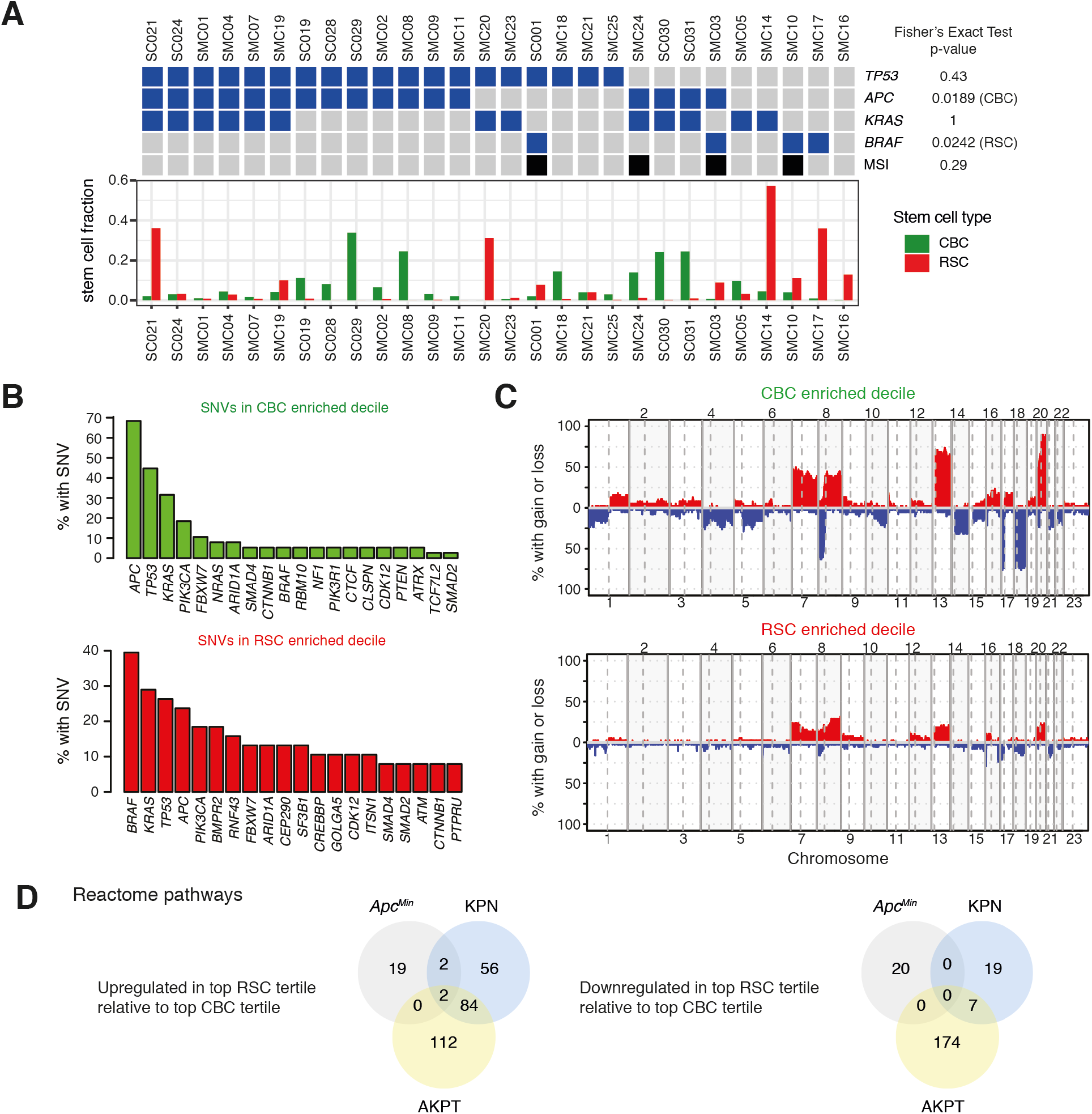
Human driver mutation landscape in CBC vs RSC enriched deciles. **A**. Fraction of epithelial cells with CBC or RSC phenotypes in single cell RNA sequencing data from human colorectal cancer patients. Mutation status of selected driver genes are marked in blue and MSI status is marked in black above each patient. Associations between CBC/RSC counts and mutation or MSI status were evaluated with a Fisher’s Exact Test (p-values listed in figure). **B**. Comparison of most prevalent single nucleotide variant mutations in TCGA tumours subdivided into CBC and RSC predominant deciles **C**. Comparison of copy number variation by chromosome number in TCGA tumours subdivided into CBC and RSC predominant deciles. **D**. Comparison of similarities and differences between upregulated or downregulated Reactome pathways in *Apc*^*Min*^, AKPT, or KPN mice from Gene Set Enrichment Analyses comparing the top tertile (highest stem cell index, most RSC) and the bottom tertile (lowest stem cell index, most CBC) of autochthonous tumors from each mouse genotype.

**Supplementary Figure 4.**
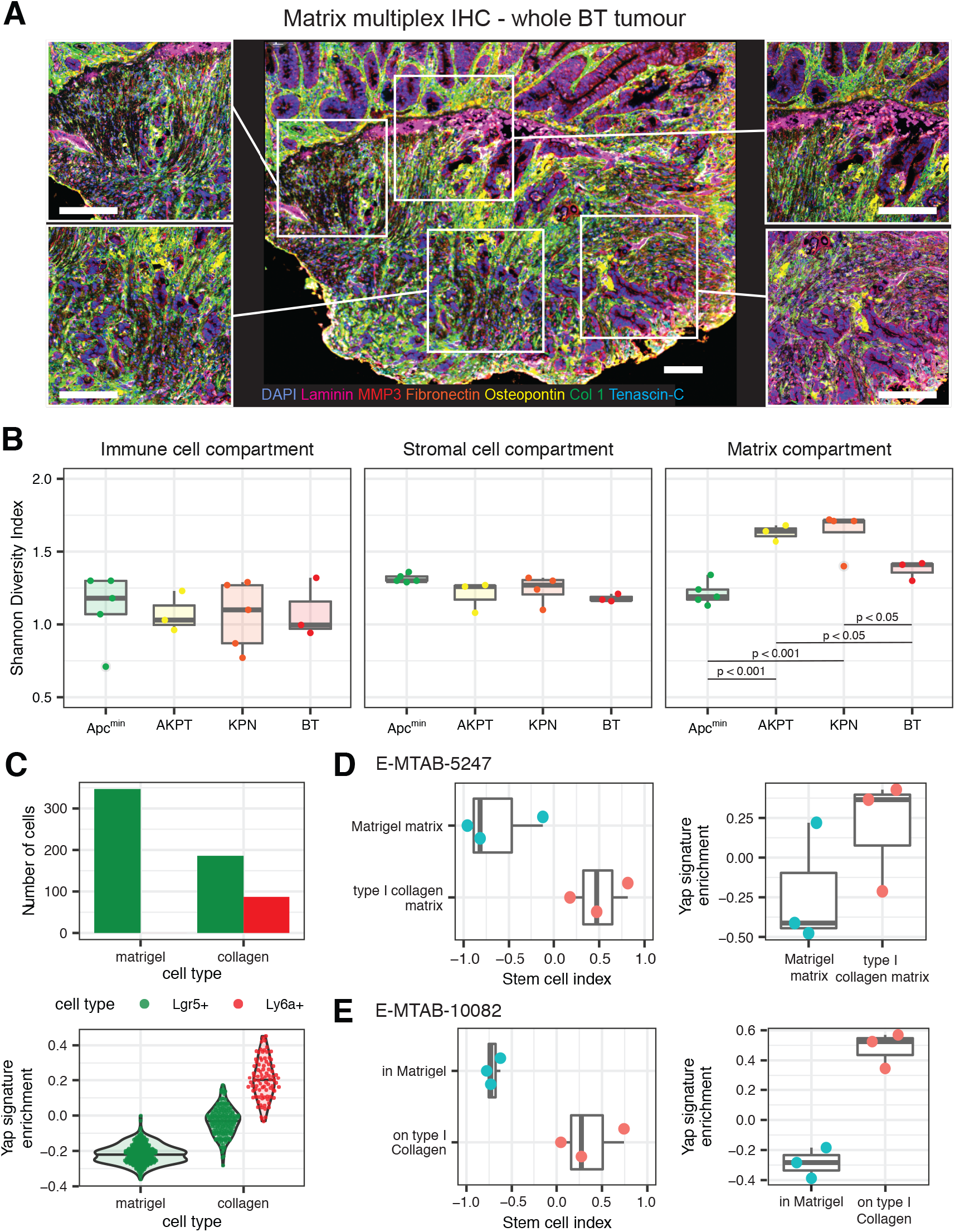
Matrix heterogeneity, diversity and impact of mechanotransduction on organoids. **A**. Topographical heterogeneity of matrix deposition detected using matrix multiplex IHC, across a representative whole tumour (BT mouse). Scale bars 100μm **B**. Cross genotype diversity of immune, stromal cell populations and matrix assessed from quantification of multiplex IHC using Shannon’s diversity index. Statistical analysis, ANOVA with Tukey post hoc test, p-values as stated. **C**. Number of *Lgr5*+ and *Ly6a*+ cells and Yap signature enrichment of single cells from organoids grown in either matrigel or collagen and analysed by single cell RNA-sequencing. **D-E**. Stem cell index and Yap signature enrichment of organoids grown in matrigel or in/on collagen analysed by bulk RNA-sequencing from **D**. E-MTAB-5247 (Yui *et al*., 2018) and **E**. E-MTAB-10082 (Ramadan *et al*., 2021).

**Supplementary Figure 5.**
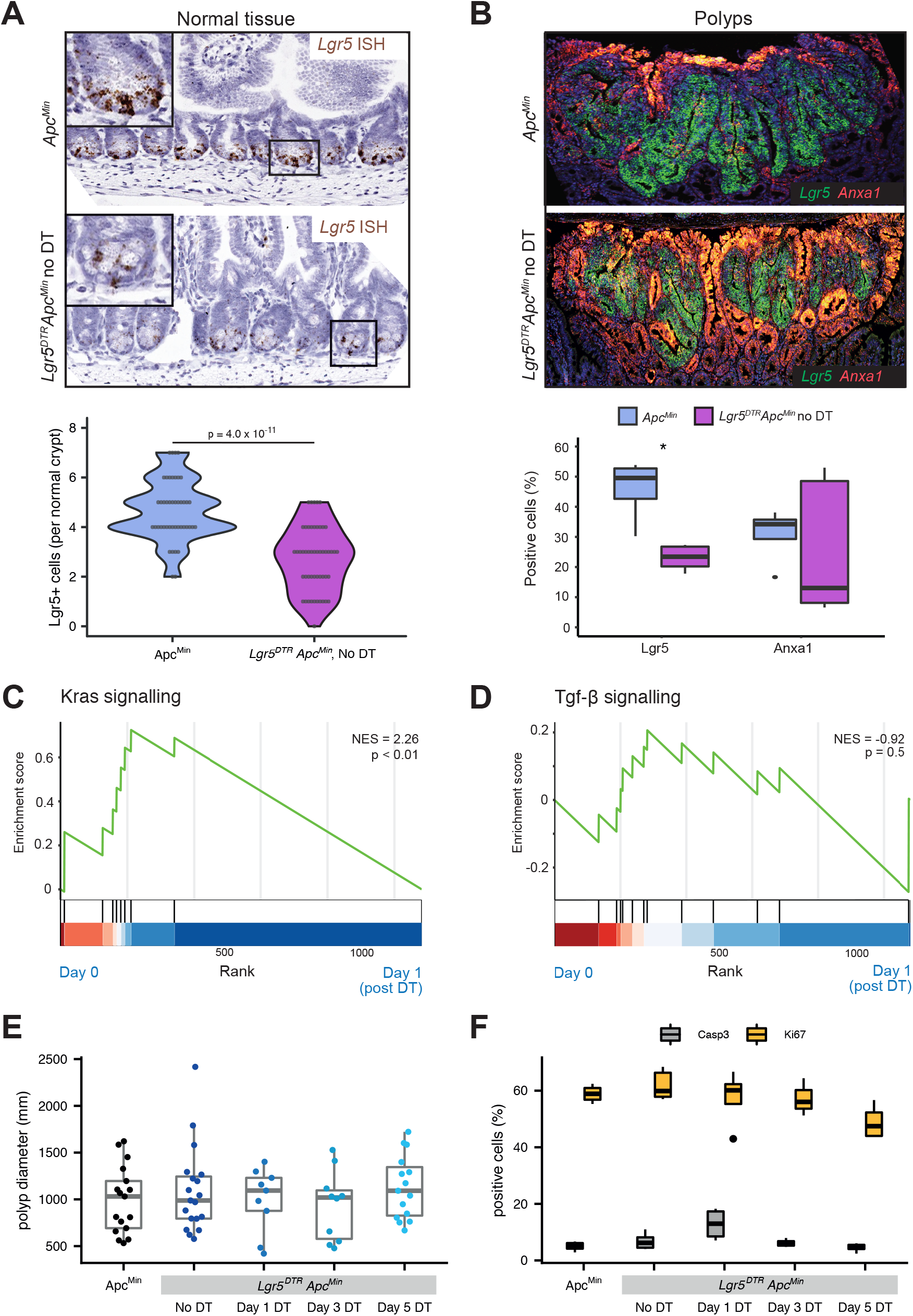
Impact of *Lgr5* hemizygosity and stem cell ablation on polyp size and proliferation in *Lgr5*^*DTR*^*;Apc*^*Min*^ mice. **A**. Chromogenic ISH and quantification showing impact of *Lgr5* hemizygosity on CBC cell count in unrecombined *Lgr5*^*DTR*^*;Apc*^*Min*^ mice. T-test, p value as stated. **B**. Dual colour fluorescent ISH and quantification showing decreased *Lgr5* expression in polyps in unrecombined *Lgr5*^*DTR*^*;Apc*^*Min*^ mice. **C-D** Gene set enrichment analysis showing enrichment of **C**. Kras signalling and **D**. lack of enrichment of Tgfβ signalling between Day 0 (unrecombined) and Day 1 (after stem cell ablation) **E**. Polyp diameter (micrometres) and **F**. cell proliferation (assessed by quantified Ki67 stain) and apoptosis (assessed by quantified caspase staining) of polyps in *Lgr5*^*DTR*^*;Apc*^*Min*^ mice before and after CBC cell ablation.

